# Bio-orthogonal chemistry-based conjugation strategy facilitates investigation of impacts of s^2^U, s^4^U, m^1^A and m^6^A guide RNA modifications on CRISPR activity

**DOI:** 10.1101/2022.06.09.495561

**Authors:** Alyssa Hoy, Ya Ying Zheng, Jia Sheng, Maksim Royzen

## Abstract

The CRISPR-Cas9 system is an important genome editing tool that holds enormous potential towards treatment of human genetic diseases. Clinical success of CRISPR technology is dependent on incorporation of modifications into the single guide RNA (sgRNA). However, chemical synthesis of modified sgRNAs, which are over 100 nucleotides in length, is difficult and low-yielding. We developed a conjugation strategy that utilized bio-orthogonal chemistry to efficiently assemble functional sgRNAs containing nucleobase modifications. The described approach entails the chemical synthesis of two shorter RNA oligonucleotides: a 31-mer containing tetrazine (Tz) group and a 70-mer modified with a *trans*-cyclooctene (TCO) moiety. The two oligonucleotides were conjugated to form functional sgRNAs. The two-component conjugation methodology was utilized to synthesize a library of sgRNAs containing nucleobase modifications such as m^1^A, m^6^A, s^2^U and s^4^U. The impacts of these RNA modifications on overall CRISPR activity was investigated *in vitro* and in Cas9-expressing HEK293T cells.

## Introduction

The CRISPR-Cas9 system is a powerful genome editing tool that profoundly revolutionized biomedical research and holds enormous potential for successful treatment of human genetic diseases. The two-component system, consisting of a single-guide RNA (sgRNA) and a DNA nuclease, Cas9, form a complex capable of targeting double stranded DNA with single nucleotide precision.(Jinek et al., 2014; Nishimasu et al., 2014) This technology has been successfully applied for genome editing in cellular level(Gaudelli et al., 2017; Wang et al., 2014), as well as *in vivo*.(Cameron et al., 2019; Finn et al., 2018; Maeder et al., 2019) In recent years, CRISPR technology has been evaluated in several human clinical trials for the treatment of blood disorders(Frangoul et al., 2021), various cancers(Lacey and Fraietta, 2020; Lu et al., 2020; Stadtmauer et al., 2020), eye disease(Ledford) and chronic infection.(Maxwell)

Virtually all FDA-approved RNA-based therapeutics contain RNA modifications.(Adams et al., 2018; Balwani et al., 2020; Jackson et al., 2020; Polack et al., 2020) Clinical success of CRISPR technology is likely dependent on well-designed incorporation of RNA modifications which can potentially improve sgRNA stability, on-target efficiency and lower off-target effects. A number of RNA modifications have already been explored to achieve these goals.(Mir et al., 2018; O’Reilly et al., 2019; Ryan et al., 2018a; Yin et al., 2018) None of the reported systems however examined sgRNA containing nucleobase modifications, which are common in nature and capable of regulating critical cellular processes.(McCown et al., 2020) This is likely due to the fact that sgRNA is over 100-nt long and solid phase synthesis (SPS) of long oligonucleotides is difficult. Furthermore, incorporation of nucleobase-modified RNA phosphoramidites during SPS exacerbates the problem, as it is typically associated with lower coupling yields. Our goal has been to develop a versatile synthetic approach that will allow to examine effects of modified nucleobases such as N^1^-methyladenosine (m^1^A), N^6^-methyladenosine m^6^A, 2-thiouridine (s^2^U) and 4-thiouridine (s^4^U) on CRISPR activity.

We were inspired by the recently reported ‘split-and-click’ strategy, illustrated in Figure 1A, which utilizes copper-catalyzed azide alkyne cycloaddition (CuAAC) chemistry to form functional sgRNAs from two smaller oligonucleotide fragments.(Taemaitree et al., 2019) This approach entails modification of the 3′-end of RNA 1 oligonucleotide with an alkyne group and modification of the 5′-end of RNA 2 oligonucleotide with an azide group. The two reacting groups are small and the resulting triazole linker has been shown to cause minimal structural perturbation to ‘clicked’ sgRNAs. One of the drawbacks of the reported approach is that the ligation chemistry is catalyzed by Cu(I), which is biologically toxic and should ideally be avoided in the synthesis of therapeutic RNAs.(Jewett and Bertozzi, 2010) Our goal was to investigate if the ‘split-and-click’ strategy could be successfully accomplished using inverse electron demand Diels-Alder (IEDDA) chemistry, which does not require Cu(I) catalysis. The bio-orthogonal IEDDA chemistry has been shown to be compatible with the intricate functional groups and structural elements of RNA.(Agustin et al., 2016; He et al., 2021; Wu et al., 2019) As illustrated in Figure 1B, our approach entails modification of the 2′-end of RNA 1 oligonucleotide with a tetrazine (Tz) group and modification of the 5′-end of RNA 2 oligonucleotide with a *trans*-cyclooctene (TCO) moiety. The two oligonucleotides can be conjugated in a biological buffer under neutral pH and without any transition metal catalyst, thus forming ‘conjugated’ sgRNA (Figure 1B). Watts and co-workers recently reported a similar bioconjugation strategy bridging two segments of sgRNA in the tetraloop region using Tz and norbornene.(Chen et al., 2022)

**Figure 1.**
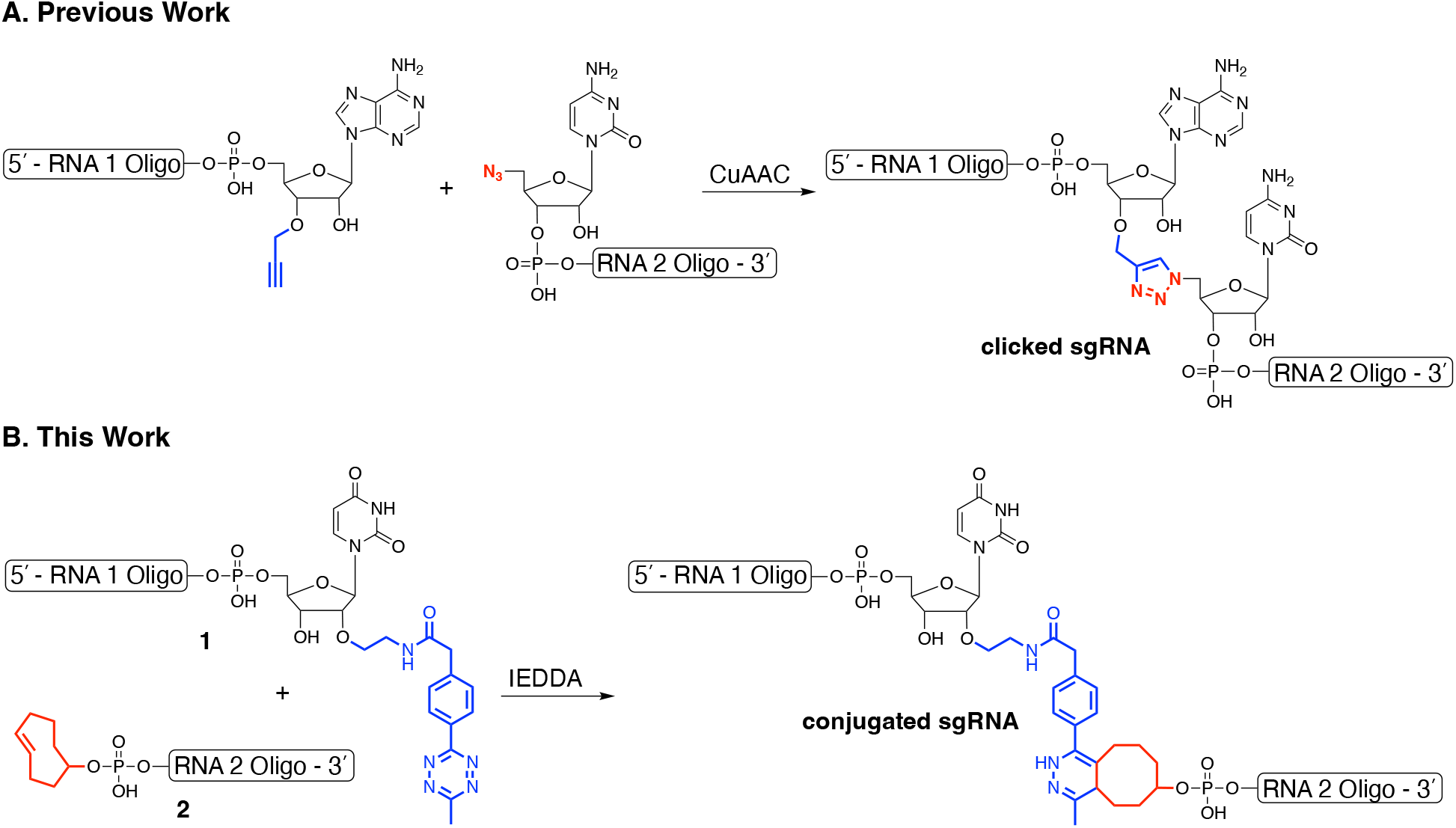
(**A**) Previously reported ‘split-and-click’ approach to form functional sgRNAs using CuAAC chemistry. (**B**) Synthesis of conjugated sgRNAs using IEDDA chemistry.

## Results

Synthesis of the two RNA components described in our strategy for making conjugated sgRNA is illustrated in Figure 2. Towards attachment of Tz to the 2′-end of RNA 1 oligonucleotide, we synthesized controlled pore glass (CPG) solid support **3** that was modified with a uridine analog containing trifluoroacetyl-protected amine group at the 2′-position.(Jin et al., 2005; Santner et al., 2014) After completion of the SPS of RNA and subsequent cleavage, deprotection and desilylation steps, the resulting RNA 1 oligonucleotide will contain an amine group that was coupled to the Tz-NHS ester **5**. Our choice of Tz was based on the reports that indicated its exceptional stability under physiological conditions.(Karver et al., 2011; Mejia Oneto et al., 2016) The Tz-modified RNA oligonucleotides were purified by preparative PAGE and characterized by ESI-MS (Figures S1-S10). As illustrated in Figure 2B, 5′-end of RNA 2 oligonucleotide was modified with TCO using previously reported TCO-phosphoramidite **7**.(Schoch et al., 2012) Compound **7** was incorporated at the final coupling step of SPS. Modified oxidation step (see SI) using *tert*-butyl hydroperoxide was used due to sensitivity of TCO to the standard SPS oxidation chemistry. The TCO-modified RNA oligonucleotides were obtained after the standard cleavage, deprotection and desilylation steps.

**Figure 2.**
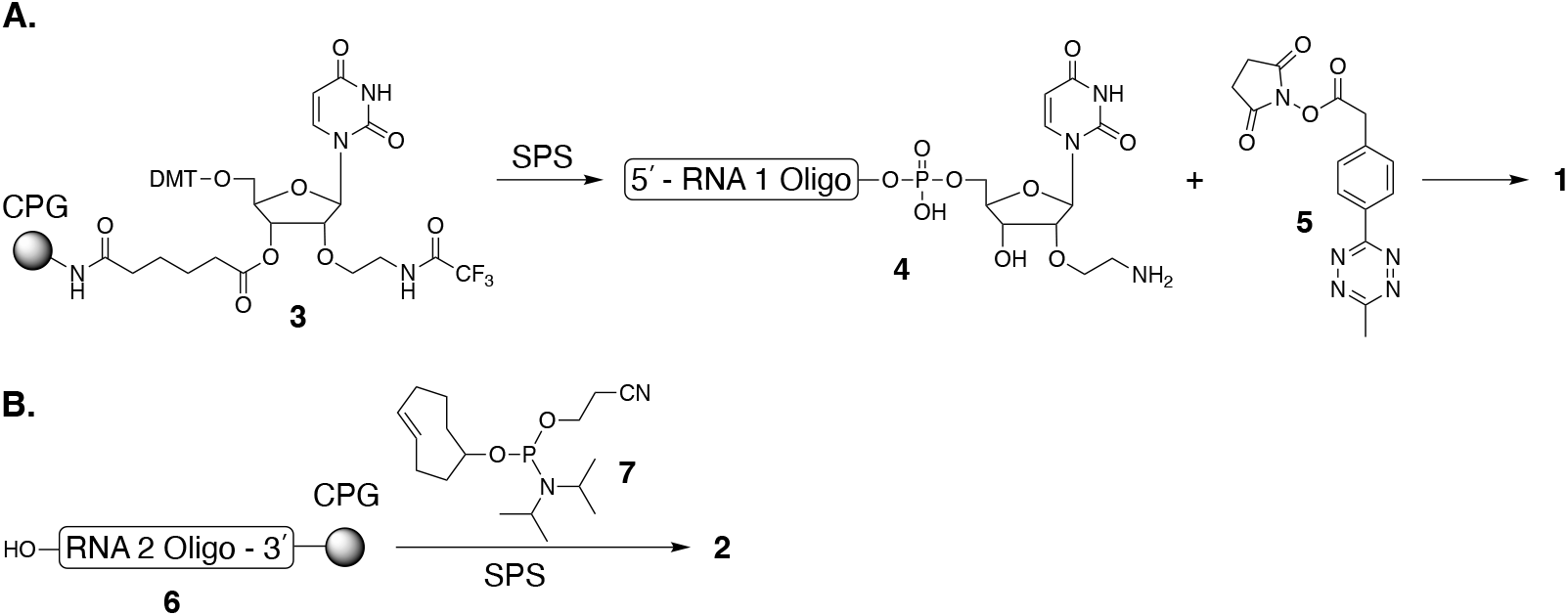
(**A**) Synthesis of the RNA oligonucleotide-modified with Tz; (**B**) Synthesis of the RNA oligonucleotide-modified with TCO.

The conjugated sgRNAs that are formed via IEDDA chemistry contain a linker that is considerably larger than the triazole linker (Figure 1A). According to experimental analysis described by Taemaitree *et al*. this could potentially be detrimental for CRISPR activity due to perturbation of native interactions between sgRNA and Cas9.(Taemaitree et al., 2019) Our goal was to find a site where longer linkers would be well-tolerated. We were inspired by Iwasaki *et al*. who reported sgRNA constructs containing fused theophylline aptamer capable of regulating CRISPR activity.(Iwasaki et al., 2020) We synthesized three sgRNAs in which the two RNA segments were conjugated together at different positions in the repeat region, as shown in Figure 3. The conjugated sgRNAs were purified by preparative PAGE, as illustrated in Figure S11. The sequence of the constructed sgRNAs has been reported to target linearized pBR322 plasmid.(Taemaitree et al., 2019)

**Figure 3.**
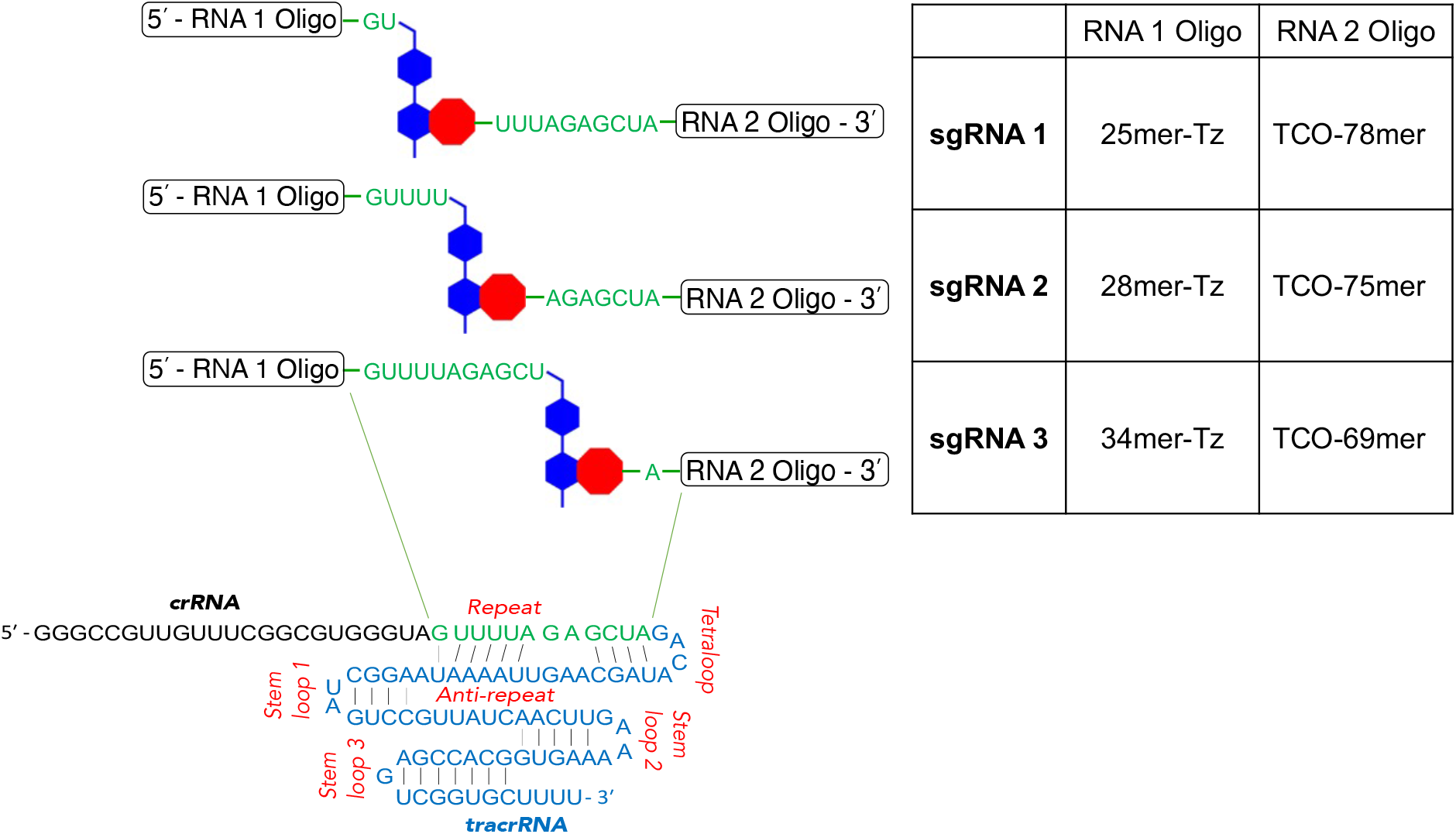
Synthesis of conjugated sgRNA constructs by ligating two RNA fragments using IEDDA chemistry.

The constructs described in Figure 3 were tested *in vitro* for their ability to carry out Cas9-mediated cleavage of linearized pBR322 plasmid. As illustrated in Figure 4, **sgRNA 1** and **sgRNA 2** were unable to facilitate DNA cleavage, presumably due to the prohibitively long linker. Fortunately, **sgRNA 3** was found to be active (lanes 6 and 7). In fact, **sgRNA 3** outperformed the unmodified sgRNA (lanes 2 and 3). These findings proved that placement of the long linker is crucial to facilitate proper RNA-Cas9 interactions that enable nuclease activity. To test the generality of our approach, we synthesized **sgRNA 4** targeting the GFP gene encoded on the linearized eGFP-N1 plasmid (Table 1, S1). The new construct was linked at the same location in the repeat sequence as **sgRNA 3**. As illustrated in Figure 5A, **sgRNA 4** (lanes 4 and 5) was also found to be active also outperforming the corresponding unmodified sgRNA (lanes 2 and 3).

**Table 1.**
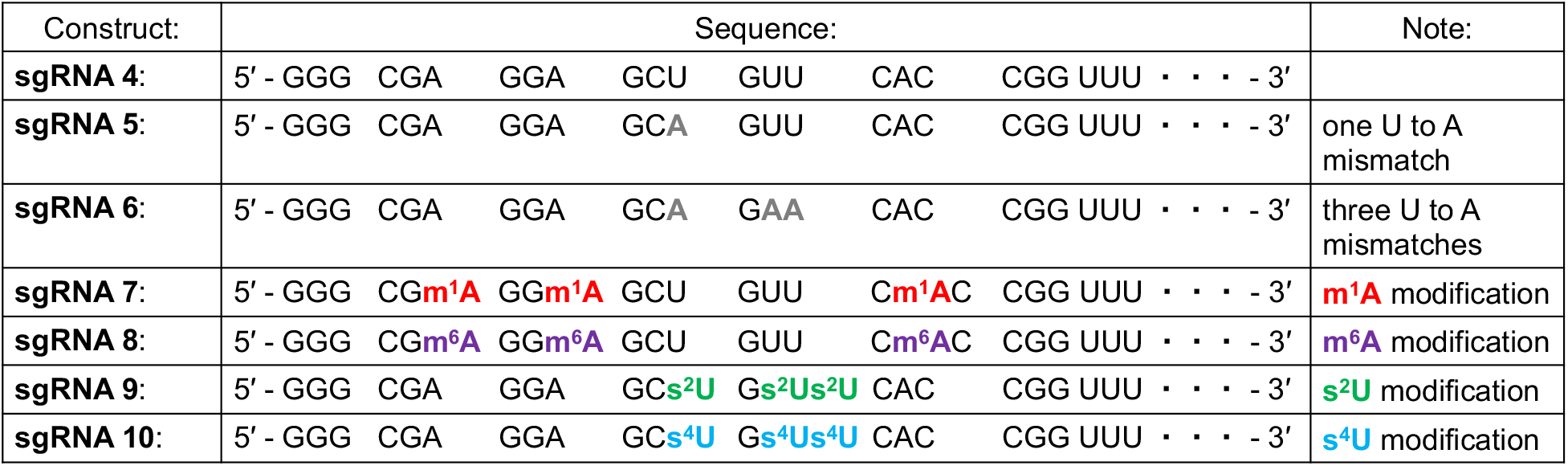
Sequences of the crRNA region of GFP-targeting sgRNAs containing nucleobase modifications.

**Figure 4.**
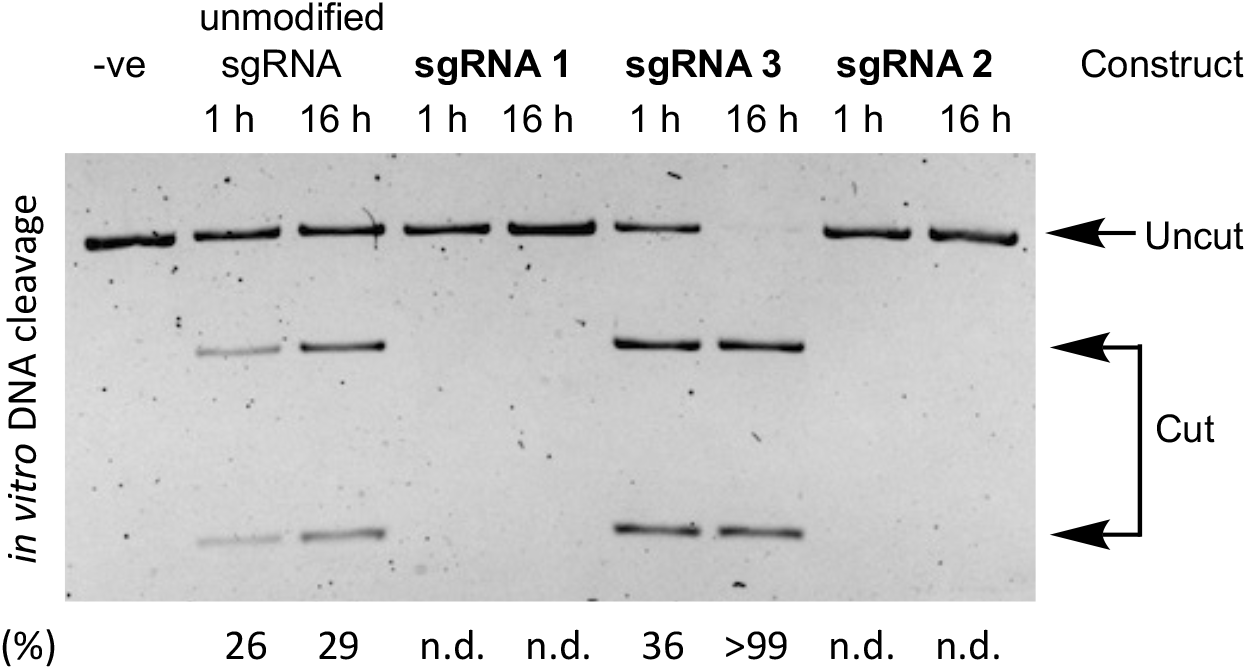
Activity of unmodified sgRNA, **sgRNA 1, sgRNA 2** and **sgRNA 3** constructs in facilitating Cas9-mediated cleavage of linearized pBR322 plasmid at 1 and 16 h time points.

**Figure 5.**
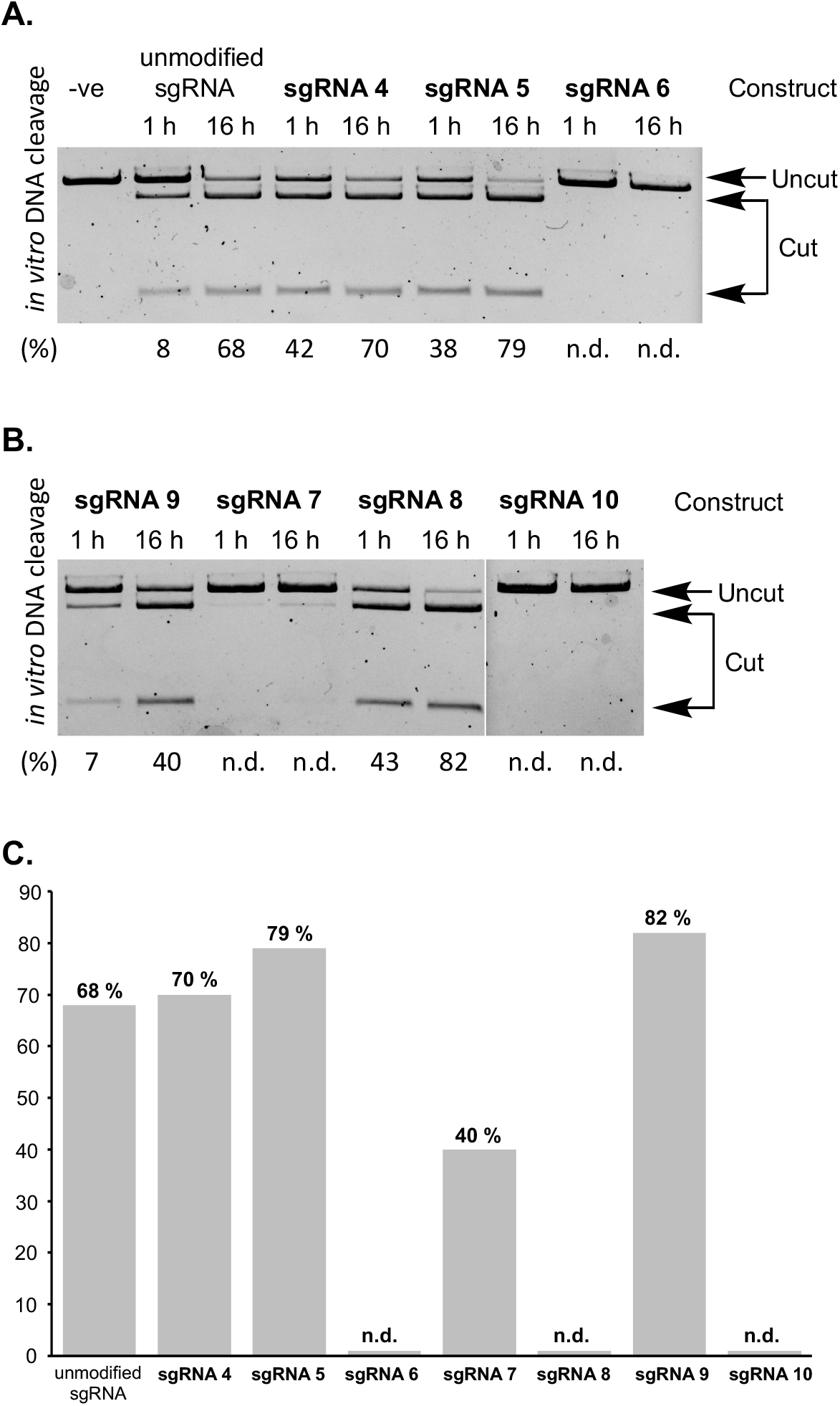
(**A**) Activity of unmodified sgRNA and **sgRNA 4, sgRNA 5** and **sgRNA 6** constructs in facilitating Cas9-mediated cleavage of linearized eGFP-N1 plasmid at 1 and 16 h time points. (**B**) Activity of sgRNAs containing m^1^A, m^6^A, s^2^U and s^4^U modifications (**sgRNA 7, sgRNA 8, sgRNA 9** and **sgRNA 10**) in facilitating Cas9-mediated cleavage of linearized eGFP-N1 plasmid at 1 and 16 h time points. (**C**) Summary of Cas9-mediated cleavage of linearized eGFP-N1 plasmid after 16 h.

Empowered by the synthetic methodology to assemble functional conjugated sgRNAs, we turned our attention towards investigation of the effects of nucleobase modifications m^1^A, m^6^A, s^2^U and s^4^U on CRISPR activity. These modifications were incorporated in the CRISPR RNA (crRNA) region (Table 1) which is responsible for recognition of complementary target DNA.(Jinek et al., 2014; Nishimasu et al., 2014) Based on previously reported studies, m^1^A, m^6^A and s^4^U will destabilize RNA-DNA duplex, while s^2^U will have stabilizing effect.(Kumar, 1997; Roost et al., 2015; Sheng et al., 2014) Our convergent approach facilitated efficient assembly of **sgRNAs 5-10** using the two-component conjugation strategy. RNA modifications, whose coupling is often associated with lower yields, were incorporated into the shorter, 31-nt long, RNA 1 oligonucleotide (Figure 1B). **sgRNAs 5**-**10** share the same 70-nt long RNA 2 oligonucleotide component (Figure 1B).

First, to get an idea of how many modifications would be necessary to observe a pronounced effect, we synthesized **sgRNA 5** and **sgRNA 6** (Table 1) that contain either one or three U to A mismatches. As illustrated in Figure 5A, a single mismatch in **sgRNA 5** does not have a pronounced impact on the nuclease activity. On the other hand, incorporation of three U to A mismatches in the **sgRNA 6** rendered it completely inactive. Therefore, to investigate the impacts of RNA modifications, we decided to synthesize **sgRNAs 7, 8, 9** and **10** containing three m^1^A, m^6^A, s^2^U and s^4^U modifications, respectively (Table 1).

As illustrated in Figure 5B, incorporation of three m^1^A modifications in **sgRNA 7** completely eliminated nuclease activity. This was an expected outcome, as methylation at the 1-position of adenosine disrupts hydrogen bonding to a complimentary thymidine. Consequently, three m^1^A residues disrupted the ability of **sgRNA 7** to recognize and bind the complimentary dsDNA. Analogous behavior was observed with **sgRNA 10**, which contains three s^4^U residues. S^4^U was reported to destabilize the duplex by 0.6 kcal/mol, which was attributed to weaker Watson–Crick base pairing due to the replacement of the hydrogen bond accepting oxygen by a weaker hydrogen bond accepting sulfur at C4.(Kumar, 1997) Meanwhile, quite unexpected results were observed with **sgRNA 8** and **sgRNA 9**. It’s been reported that s^2^U modification lead to increased duplex stability by about 1 kcal/mol.(Sheng et al., 2014; Testa et al., 1999) The increased stability has been attributed to enhanced stacking interactions and higher acidity of the N-3 proton and thus increased strength of its hydrogen bond.(Sheng et al., 2014; Testa et al., 1999) However, incorporation of three s^2^U residues in **sgRNA 9** led to lower CRISPR activity relative to **sgRNA 4**, which contains canonical nucleobases. On the other hand, **sgRNA 8**, containing three m^6^A residues exhibited higher nuclease activity than **sgRNA 4**. This finding was also unexpected, as N^6^-methylation is known to have destabilizing effect in base-paired regions of RNA by 0.5-1.7 kcal/mol. The N^6^-methyl group has been predicted to be in a high-energy anti conformation, with the methyl group oriented into the major groove of the RNA, where the amino proton would normally be.(Roost et al., 2015)

Ability of conjugated sgRNAs, containing m^6^A and s^2^U modifications, to enable Cas9 nuclease activity was tested in GFP and Cas9-expressing HEK293T cells.(Yin et al., 2018) The cells were transfected for 24 h with the GFP-targeting **sgRNA 4, sgRNA 8** and **sgRNA 9**, as well as unmodified sgRNA, as a positive control. The cells were analyzed by flow cytometry 48 h post transfection to investigate the expression of GFP. As illustrated in Figure S12, the untransfected cells show two sub-populations of GFP expressing cells. The subpopulation with higher GFP fluorescence corresponds to 4.16 ± 0.44 % of total cells, while the subpopulation with lower GFP signal corresponds to 78.93 ± 0.25 % of total cells. Transfection with unmodified sgRNA almost completely eliminated the subpopulation of cells with higher GFP fluorescence (0.25 ± 0.14 %), while also decreasing the subpopulation of cells with lower GFP signal to 66.37 ± 3.69 %. Analogous behavior was observed with the cells transfected with **sgRNA 4** (higher GFP subpopulation 0.20 ± 0.05 %; lower GFP subpopulation 67.07 ± 1.18 %). This indicated that **sgRNA4** is functional inside the cells to enable nuclease activity that is on par with the native system. Contrary to the prediction from the *in vitro* experiments, **sgRNA 8** and **sgRNA 9** were found to have lower CRISPR activity inside the cells. The flow cytometry analysis illustrated in Figure S13, indicates that the cells transfected with **sgRNA 8** had a higher GFP-expressing subpopulation of 2.12 ± 0.07 % and lower GFP-expressing subpopulation of 74.07 ± 0.60 %. Similarly, the cells transfected with **sgRNA 9** had a higher GFP-expressing subpopulation of 2.31 ± 0.11 % and lower GFP-expressing subpopulation of 69.80 ± 0.85 %. These data are summarized in Figure S13.

## Discussion

The CRISPR-Cas9 system is a powerful genome editing tool whose clinical translation will likely require thorough structural optimization. Cas9-guided nuclease activity is dependent on the interactions between the enzyme and sgRNA, as well as the subsequent recognition and association between this Cas9-sgRNA complex and target dsDNA. This multi-step process has been demonstrated by structural studies to involve a cascade of conformational changes within the RNA-enzyme-DNA system.(Jiang and Doudna, 2017; Sternberg et al., 2015) In recent years, a number of different strategies have evolved to manipulate and optimize each of the three components and improve the overall CRISPR efficiency and specificity of DNA cleavage. For example, new and improved Cas9 variants have been reported, such as eSpCas9, SpCas9-HF1, and HypaCas9.(Chen et al., 2017; Kleinstiver et al., 2016; Slaymaker et al., 2016) In comparison, the strategy of changing the sequence, stability or conformation of guide RNA requires relatively less effort than new enzyme engineering. It has been shown that chemical modifications on guide RNA such as 2′-deoxy, 2′-F, 2′-O-methyl, LNA and phosphorothioated backbone can improve nuclease activity and gene editing efficiency probably due to both enhanced interactions within the CRISPR system and increased biostability of RNA.(Hendel et al., 2015; Rahdar et al., 2015; Rueda et al., 2017; Ryan et al., 2018b; Yin et al., 2017)

However, balancing and optimizing the Cas9 cleavage efficiency and on-target specificity remains a big challenge for gene editing and requires more systematic investigation. From this perspective, using naturally or artificially modified nucleobases to further adjust the interactions of crRNA, Cas9 and target DNA represents a promising approach. There are many base modifications that are currently available to fine-tune the stability and specificity of RNA binding with both Cas9 and DNA. Due to the intrinsic synthetic challenges of making long base-modified RNA strands, such methods have not been well explored. Our metal-free IEDDA chemistry-based bio-orthogonal method improves the RNA synthesis yield and facilitates construction of a model platform to thoroughly study the overall DNA cleavage effects of all the base modifications with available phosphoramidite building blocks.

In the current work, we demonstrated that the TCO-Tz linkages can be compatible within the CRISPR system if placed in the optimized position in sgRNA. We selected different numbers of m^1^A, m^6^A, s^4^U and s^2^U, which are all natural RNA modifications, to decorate the sgRNA as the proof-of-concept study focusing on DNA cleavage efficiency. Since m^1^A and m^6^A destabilize RNA-DNA duplex, while s^2^U and s^4^U have stabilizing effect, we set out to test correlation with the overall CRISPR efficiency.

It turns out three s^4^U or m^1^A residues completely inhibit the DNA cleavage activity probably due to the resulting unstable crRNA-DNA duplex. In comparison, three m^6^A significantly increase the cleavage activity, which is contradictory to its destabilization effect on RNA-DNA. Similarly, three s^2^U residues, which are expected to largely enhance the duplex stability, have slightly negative impact on the cleavage activity. These results indicate that there might be no direct correlations between the stabilization effect and CRISPR activity. Although more systematic structural and biophysical insights are required to investigate the impacts of these modifications on Cas9 interactions, most likely the enzyme recognition of crRNA has been changed with these modifications. In addition, for the modifications such as m^6^A that can increase the DNA cleavage efficiency, the cleavage specificity or off-target effect remain to be investigated. Nonetheless, this work provides a useful platform to test more chemical modifications and identify some useful candidates to further increase both CRISPR efficiency and precision.

## Acknowledgments

We gratefully acknowledge financial support by the National Institutes of Health of the United States (Grant R21HG012257 to M.R.) and the National Science Foundation (Grant CHE-1664577 to M.R.), (Grants CHE-1845486 to J.S.). This work was also supported, in part, by the National Institutes of Health of the United States Grant R15GM12811501.

## Author Contributions

A.H. carried out chemical synthesis of RNA phosphoramidites and SPS of RNA containing m^1^A and m^6^A modifications. A.H. performed *in vitro* CRISPR experiments. Y.Y.Z. carried out SPS of RNA containing s^2^U and s^4^U modifications. Y.Y.Z. performed in-cell CRISPR experiments and flow cytometry. M. R. and J. S. conceptualized the project, provided student supervision and funding for the project. M. R. and J. S. wrote the manuscript with input from all authors. All authors approved the final manuscript.

## Supplemental Information

Supplementary information can be found at:

