## Supporting Information for "Bio-orthogonal chemistry-based conjugation strategy facilitates investigation of impacts of s^2^U, s^4^U, m^1^A and m^6^A guide RNA modifications on CRISPR activity"



| **Table of Contents** | **Figure and Scheme titles** | **Page** |
|---|---|---|
| Materials and Methods | | S2-S4 |
| Sequences of the reported RNAs | | S5-S9 |
| Table S1 | ESI-MS analysis of the reported RNA oligonucleotides. | S10 |
| Figure S1 | Deconvoluted ESI-MS spectrum of the 25mer-Tz that was used to form **sgRNA1**. | S11 |
| Figure S2 | Deconvoluted ESI-MS spectrum of the 28mer-Tz that was used to form **sgRNA2**. | S11 |
| Figure S3 | Deconvoluted ESI-MS spectrum of the 34mer-Tz that was used to form **sgRNA3**. | S12 |
| Figure S4 | Deconvoluted ESI-MS spectrum of the 31mer-Tz that was used to form **sgRNA4**. | S12 |
| Figure S5 | Deconvoluted ESI-MS spectrum of the 31mer-Tz that was used to form **sgRNA5**. | S13 |
| Figure S6 | Deconvoluted ESI-MS spectrum of the 31mer-Tz that was used to form **sgRNA6**. | S13 |
| Figure S7 | Deconvoluted ESI-MS spectrum of the 31mer-Tz that was used to form **sgRNA7**. | S14 |
| Figure S8 | Deconvoluted ESI-MS spectrum of the 31mer-Tz that was used to form **sgRNA8**. | S14 |
| Figure S9 | Deconvoluted ESI-MS spectrum of the 31mer-Tz that was used to form **sgRNA9**. | S15 |
| Figure S10 | Deconvoluted ESI-MS spectrum of the 31mer-Tz that was used to form **sgRNA10**. | S15 |
| Figure S11 | PAGE analysis of conjugation of 25mer-Tz and TCO-78mer. | S16 |
| Figure S12 | Flow cytometry analysis of GFP expression of GFP and Cas9-expressing HEK293T transfected with native sgRNA and **sgRNA4**. | S17 |
| Figure S13 | Flow cytometry analysis of GFP expression of GFP and Cas9-expressing HEK293T transfected with **sgRNA8** and **sgRNA9**. | S18 |
| Figure S14 | Flow cytometry analysis of GFP expression of GFP and Cas9-expressing HEK293T transfected with unmodified sgRNA, **sgRNA 4**, **sgRNA8** and **sgRNA9**. | S19 |
| Synthetic Procedures | Flow cytometry analysis of CRISPR experiments. | S20-S21 |
| NMR Spectra | | S22-S24 |

### Materials and Methods

All oligonucleotide solid phase syntheses were done on a 1.0 µmol scale using the Oligo-800 synthesizer (Azco Biotech, Oceanside, CA, USA). Solid phase syntheses were performed on control-pore glass (CPG-1000) purchased from Glen Research (Sterling, VA, USA). Other oligonucleotide solid phase synthesis reagents were obtained from ChemGenes Corporation (Wilmington, MA, USA). Phosphoramidites (TBDMS as the 2′-OH protecting group): rA was N-Bz protected, rC was N-Ac protected and rG was N-iBu protected. A, C, G, U phosphoramidites were dissolved in anhydrous acetonitrile (0.07 M) directly before use. $m^1A$, $m^6A$, $s^2U$ and $s^4U$ phosphoramidites were dissolved in anhydrous acetonitrile (0.15 M) directly before use. Coupling step was done using 5-ethylthio-1H-tetrazole solution (0.25 M) in acetonitrile for 12 min. 5′-detritylation step was done using 3% trichloroacetic acid in $CH_2Cl_2$. Oxidation step was done using $I_2$ (0.02 M) in THF/pyridine/$H_2O$ solution. CPG modifications were carried out using native amino lcaa CPG 1000 Å, purchased from ChemGenes (Wilmington, MA, USA), Cat.# N-5100-10).

For gel electrophoresis, 10X Tris/Borate/EDTA (TBE) buffer was purchased from Fisher Scientific Company L.L.C. (Waltham, MA, USA) and used with proper dilution. 30% Arcylamide/Bis-arcylamide solution (29:1) was purchased from Bio-Rad Laboratories, Inc. (Hercules, CA, USA).

Chromatographic purifications of synthetic materials were conducted using SiliaSphere™ spherical silica gel with an average particle and pore size of 5 µm and 60 Å, respectively (Silicycle Inc, QC, Canada). Thin layer chromatography (TLC) was performed on SiliaPlate™ silica gel TLC plates with 250 µm thickness (Silicycle Inc, QC, Canada). Flash chromatography was performed using Biotage Isolara One instrument (Biotage Sweden AB, Uppsala, Sweden). Preparative TLC was performed using SiliaPlate™ silica gel TLC plates with 1000 µm thickness. $^{1}H$, $^{13}C$ and $^{31}P$ NMR spectroscopy was performed on a Bruker NMR at 500 MHz ($^{1}H$) and 126 MHz ($^{13}C$) and 121 MHz ($^{31}P$). All $^{13}C$ NMR spectra were proton decoupled. High resolution ESI-MS spectra of small molecules was acquired using Agilent Technologies 6530 Q-TOF instrument. ESI-MS analysis of RNA oligonucleotides was carried out by Novatia, LLC (Novatia, LLC, Newtown, PA).

### Synthesis of RNA oligonucleotides containing TCO

The phosphoramidite **7,** containing TCO, was coupled during the final synthetic cycle of SPS of RNA. The subsequent oxidation step was performed using 1 M solution of t-BuOOH in $CH_2Cl_2$. After completion of SPS, CPG beads were treated with concentrated aqueous ammonia in a sealed vial for 2 h at 55 °C. The ammonia was removed in vacuo (SpeedVac). Cleaved oligonucleotides were redissolved in mixture of anhydrous DMSO (100 µL) and triethylamine trihydrofluoride (125 µL) and heated for 2.5 h at 65 °C. After subsequent cooling to rt, sodium acetate buffer (3 M, pH 5.2, 25 µL) and ethanol (1 mL) were added, and the RNA was stored overnight at −20 °C. The RNA was then pelleted by centrifugation (14,000 × *g*, 17 min, 4 °C), the supernatant was discarded, and the pellet was washed with 75% ethanol (1 mL). The pellet was then dried in vacuo, dissolved in water.

### Assessment of TCO coupling to RNA oligonucleotides

CPG beads containing RNA 2 oligo-TCO (5 mg) were mixed with Tz-DMT for 1 h at rt. Synthesis of Tz-DMT was previously described [He, M.; Wu, X.; Mao, S.; Haruehanroengra, P.; Khan, I.; Sheng, J. Royzen, M. *Chem. Comm.* **2021**, *57*, 4263-4266]. After the reaction, the beads were thoroughly washed with DMF and $CH_2Cl_2$. The CPG beads were treated with 10 mL deblocking solution (3% trichloroacetic acid in $CH_2Cl_2$). Absorbance at 504 nm was used to calculate the amount of cleaved DMT group (ε = 76 mL $cm^{-1}$ $\mu mol^{-1}$). The amount of coupled TCO was calculated based on the assumption of nearly quantitative reaction between RNA 2 oligo-TCO and Tz-DMT.

### Purification of RNA oligonucleotides by preparative PAGE

Oligonucleotides were mixed with formamide dye (50% v/v) and loaded onto a denaturing 10-20% polyacrylamide gel (1× TBE buffer containing 8 M urea and separated at 500 V for 1-2 h. RNA bands were visualized under UV, excised, crushed, soaked in buffer (ammonium acetate 500 mM, $Mg(OAc)_2$ 10 mM, EDTA 2mM) 800µL) for 16 h at rt with vigorous shaking. Buffer volume was reduced to 50 µL by butanol extraction (5x500 µL). Cold ethanol (1000 µL) was added and RNA was precipitated at -20 °C overnight.

### Cas9 *in vitro* cleavage assay.

pBR322 plasmid DNA (0.35 µM, 1.13 µL, NEB, N3033S) was diluted with water (16.87 µL) and NEB buffer 3.1 (10x, 2 µL). The plasmid was linearized directly prior to CRISPR experiments using PvuII (10 U/µL, 1 µL, NEB, R0151S) for 1 h at 37 °C. In analogous fashion eGFP-N1 plasmid was linearized with DraIII-HF (10 U/µL, 1 µL, NEB, R3510L). For the Cas9-mediated DNA cleavage assay, sgRNA (300 nM, 5 µL), Cas9 (1 µM, 0.3 µL, NEB, M0386S), Cas9 buffer (10x, 1 µL, NEB), linearised plasmid (20 nM, 1.5 µL) and $H_2O$ (2.2 µL) were mixed together (final volume = 10 µL) and incubated for either 1 or 16 h at 37 °C. CRISPR experiments were terminated by the addition of proteinase K (20 mg/mL, 0.5 µL) for 1 h at 37 °C. The reaction (1 µL) was mixed with blue loading buffer (6x, 2 µL, NEB, B7703S) and loaded on a 1% agarose stained with ethidium bromide (1x TBE running buffer).

### Evaluation of in-cell CRISPR activity

CRISPR-Cas9 experiments, were carried out following the procedure reported by Yin, H. *et al.* [*Nat. Chem. Biol.* **2018**, *14*, 311-316]. HEK293T cells were infected by lentiviral particles to stably express EF1a-GFP-PGK-Puro (Addgene; 26777) and EFs-spCas9-Blast (Addgene; 52962). The cells were grown to 70-90% confluence in DMEM, containing 10% FBS. The cells were transfected with GFP-targeting sgRNAs (30 nM each, final concentration) using Lipofectamine (Thermo Fisher Scientific) for 24 h. The cells were further grown in DMEM for 48 h post-transfection. The cells were fixed with 2% paraformaldehyde in PBS and GFP expression was analyzed by flow cytometry. Data from $10^6$ cells were acquired using a FACS Aria III cell sorter equipped with a 488 nm/blue coherent sapphire solid-state laser, 20 mW (BD Biosciences, San Jose, CA, USA). Data analyses were carried out using FlowJo software (Ashland, OR, USA), according to manufacturer's instructions. Parameters, such as MFI and the percentages of specific populations were quantified by histogram analysis.

### Modification of RNA 1 Oligo with Tz

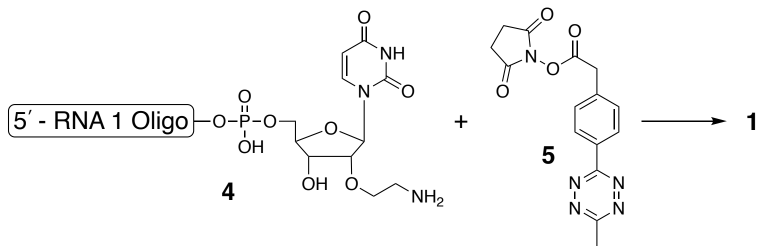


RNA 1 oligonucleotide (263 μM) was dissolved in borate buffer (250 μL, pH = 9.4). Compound **5** was dissolved in DMF (400μL, 50 mM) and added to the RNA. The reaction mixture was vortexed and agitated at 1000 rpm at rt for 8 h. The RNA 1 oligonucleotide **1** was purified from small molecules using Amicon Ultra 3K column (MilliporeSigma, cat# UFC500396) and washed with MQ $H_2O$ (3x 300 μL).

### Synthesis of conjugated sgRNA using IEDDA chemistry

RNA 1 oligo **1** containing Tz (250 μmol) and RNA 2 oligo **2** containing TCO (120 μmol) were mixed in a NaCl solution (0.2 M, 7.9 μL), containing EDTA (0.5 M, 0.2 μL) and HEPES (0.2 M, pH 7.5, 2 μL). The reaction mixture was agitated at 1000 rpm at rt for 16 h. The conjugated sgRNA was purified by preparative PAGE.

**Sequences of the reported RNAs:**

pBR322 plasmid targeting **sgRNA 1**

5′ - (rG)(rG)(rG)(rC)(rG)(rC)(rU)(rU)(rG)(rU)(rU)(rU)(rC)(rG)(rG)(rC)(rG)(rU)(rG)(rG)(rG)(rU)(rA)(rG)-**Tz-TCO**-(rU)(rU)(rU)(rU)(rA)(rG)(rA)(rG)(rC)(rU)(rA)(rG)(rA)(rC)(rA)(rU)(rA)(rG)(rC)(rA)(rA) (rG)(rU)(rU)(rA)(rA)(rA)(rA)(rU)(rA)(rA)(rG)(rG)(rC)(rU)(rA)(rG)(rU)(rC)(rC)(rG)(rU)(rU)(rA)(rU)(rC)(rA)(rA)(rC)(rU)(rU)(rG)(rA)(rA)(rA)(rA)(rA)(rG)(rU)(rG)(rG)(rC)(rA)(rC)(rC)(rG)(rA)(rG)(rU)(rC)(rG)(rG)(rU)(rG)(rC)(rU)(rU)(rU)(rU) - 3′

Synthesized by conjugation of:

5′ - (rG)(rG)(rG)(rC)(rG)(rC)(rU)(rU)(rG)(rU)(rU)(rU)(rC)(rG)(rG)(rC)(rG)(rU)(rG)(rG)(rG)(rU)(rA)(rG)-**Tz** - 3′

and

5′ - **TCO**-(rU)(rU)(rU)(rU)(rA)(rG)(rA)(rG)(rC)(rU)(rA)(rG)(rA)(rC)(rA)(rU)(rA)(rG)(rC)(rA)(rA) (rG)(rU)(rU)(rA)(rA)(rA)(rA)(rU)(rA)(rA)(rG)(rG)(rC)(rU)(rA)(rG)(rU)(rC)(rC)(rG)(rU)(rU)(rA)(rU)(rC)(rA)(rA)(rC)(rU)(rU)(rG)(rA)(rA)(rA)(rA)(rA)(rG)(rU)(rG)(rG)(rC)(rA)(rC)(rC)(rG)(rA)(rG)(rU)(rC)(rG)(rG)(rU)(rG)(rC)(rU)(rU)(rU)(rU) - 3′

pBR322 plasmid targeting **sgRNA 2**

5′ - (rG)(rG)(rG)(rC)(rG)(rC)(rU)(rU)(rG)(rU)(rU)(rU)(rC)(rG)(rG)(rC)(rG)(rU)(rG)(rG)(rG)(rU)(rA)(rG)(rU)(rU)(rU)(rU)-**Tz-TCO**-(rA)(rG)(rA)(rG)(rC)(rU)(rA)(rG)(rA)(rC)(rA)(rU)(rA)(rG)(rC)(rA)(rA) (rG)(rU)(rU)(rA)(rA)(rA)(rA)(rU)(rA)(rA)(rG)(rG)(rC)(rU)(rA)(rG)(rU)(rC)(rC)(rG)(rU)(rU)(rA)(rU)(rC)(rA)(rA)(rC)(rU)(rU)(rG)(rA)(rA)(rA)(rA)(rA)(rG)(rU)(rG)(rG)(rC)(rA)(rC)(rC)(rG)(rA)(rG)(rU)(rC)(rG)(rG)(rU)(rG)(rC)(rU)(rU)(rU)(rU) - 3′

Synthesized by conjugation of:

5′ - (rG)(rG)(rG)(rC)(rG)(rC)(rU)(rU)(rG)(rU)(rU)(rU)(rC)(rG)(rG)(rC)(rG)(rU)(rG)(rG)(rG)(rU)(rA)(rG)(rU)(rU)(rU)(rU)-**Tz** - 3′

and

5′ - **TCO**-(rA)(rG)(rA)(rG)(rC)(rU)(rA)(rG)(rA)(rC)(rA)(rU)(rA)(rG)(rC)(rA)(rA) (rG)(rU)(rU)(rA)(rA)(rA)(rA)(rU)(rA)(rA)(rG)(rG)(rC)(rU)(rA)(rG)(rU)(rC)(rC)(rG)(rU)(rU)(rA)(rU)(rC)(rA)(rA)(rC)(rU)(rU)(rG)(rA)(rA)(rA)(rA)(rA)(rG)(rU)(rG)(rG)(rC)(rA)(rC)(rC)(rG)(rA)(rG)(rU)(rC)(rG)(rG)(rU)(rG)(rC)(rU)(rU)(rU)(rU) - 3′

pBR322 plasmid targeting **sgRNA 3**

5′ - (rG)(rG)(rG)(rC)(rG)(rC)(rU)(rU)(rG)(rU)(rU)(rU)(rC)(rG)(rG)(rC)(rG)(rU)(rG)(rG)(rG)(rU)(rA)(rG)(rU)(rU)(rU)(rU)(rA)(rG)(rA)(rG)(rC)(rU)-**Tz-TCO**-(rA)(rG)(rA)(rC)(rA)(rU)(rA)(rG)(rC)(rA)(rA) (rG)(rU)(rU)(rA)(rA)(rA)(rA)(rU)(rA)(rA)(rG)(rG)(rC)(rU)(rA)(rG)(rU)(rC)(rC)(rG)(rU)(rU)(rA)(rU)(rC)(rA)(rA)(rC)(rU)(rU)(rG)(rA)(rA)(rA)(rA)(rA)(rG)(rU)(rG)(rG)(rC)(rA)(rC)(rC)(rG)(rA)(rG)(rU)(rC)(rG)(rG)(rU)(rG)(rC)(rU)(rU)(rU)(rU) - 3′

Synthesized by conjugation of:

5′ - (rG)(rG)(rG)(rC)(rG)(rC)(rU)(rU)(rG)(rU)(rU)(rU)(rC)(rG)(rG)(rC)(rG)(rU)(rG)(rG)(rG)(rU)(rA)(rG)(rU)(rU)(rU)(rU)(rA)(rG)(rA)(rG)(rC)(rU)-**Tz** - 3′

and

5′ - **TCO**-(rA)(rG)(rA)(rC)(rA)(rU)(rA)(rG)(rC)(rA)(rA) (rG)(rU)(rU)(rA)(rA)(rA)(rA)(rU)(rA)(rA)(rG)(rG)(rC)(rU)(rA)(rG)(rU)(rC)(rC)(rG)(rU)(rU)(rA)(rU)(rC)(rA)(rA)(rC)(rU)(rU)(rG)(rA)(rA)(rA)(rA)(rA)(rG)(rU)(rG)(rG)(rC)(rA)(rC)(rC)(rG)(rA)(rG)(rU)(rC)(rG)(rG)(rU)(rG)(rC)(rU)(rU)(rU)(rU) - 3′

eGFP-N1plasmid targeting **sgRNA 4**

5′ - (rG)(rG)(rG)(rC)(rG)(rA)(rG)(rG)(rA)(rG)(rC)(rU)(rG)(rU)(rU)(rC)(rA)(rC)(rC)(rG)(rG)(rU)(rU)(rU)(rU)(rA)(rG)(rA)(rG)(rC)(rU)-**Tz-TCO**-(rA)(rG)(rA)(rA)(rA)(rU)(rA)(rG)(rC)(rA)(rA) (rG)(rU)(rU)(rA)(rA)(rA)(rA)(rU)(rA)(rA)(rG)(rG)(rC)(rU)(rA)(rG)(rU)(rC)(rC)(rG)(rU)(rU)(rA)(rU)(rC)(rA)(rA)(rC)(rU)(rU)(rG)(rA)(rA)(rA)(rA)(rA)(rG)(rU)(rG)(rG)(rC)(rA)(rC)(rC)(rG)(rA)(rG)(rU)(rC)(rG)(rG)(rU)(rG)(rC)(rU)(rU)(rU)(rU)(rU) - 3′

Synthesized by conjugation of:

5′ - (rG)(rG)(rG)(rC)(rG)(rA)(rG)(rG)(rA)(rG)(rC)(rU)(rG)(rU)(rU)(rC)(rA)(rC)(rC)(rG)(rG)(rU)(rU)(rU)(rU)(rA)(rG)(rA)(rG)(rC)(rU)-**Tz** - 3′

and

5′ - **TCO**-(rA)(rG)(rA)(rA)(rA)(rU)(rA)(rG)(rC)(rA)(rA) (rG)(rU)(rU)(rA)(rA)(rA)(rA)(rU)(rA)(rA)(rG)(rG)(rC)(rU)(rA)(rG)(rU)(rC)(rC)(rG)(rU)(rU)(rA)(rU)(rC)(rA)(rA)(rC)(rU)(rU)(rG)(rA)(rA)(rA)(rA)(rA)(rG)(rU)(rG)(rG)(rC)(rA)(rC)(rC)(rG)(rA)(rG)(rU)(rC)(rG)(rG)(rU)(rG)(rC)(rU)(rU)(rU)(rU)(rU) - 3′

eGFP-N1plasmid targeting **sgRNA 5** containing one U to A mismatch

5′ - (rG)(rG)(rG)(rC)(rG)(rA)(rG)(rG)(rA)(rG)(rC)(**rA**)(rG)(rU)(rU)(rC)(rA)(rC)(rC)
(rG)(rG)(rU)(rU)(rU)(rU)(rA)(rG)(rA)(rG)(rC)(rU)-**Tz-TCO**-(rA)(rG)(rA)(rA)(rA)(rU)(rA)
(rG)(rC)(rA)(rA) (rG)(rU)(rU)(rA)(rA)(rA)(rA)(rU)(rA)(rA)(rG)(rG)(rC)(rU)(rA)(rG)(rU)
(rC)(rC)(rG)(rU)(rU)(rA)(rU)(rC)(rA)(rA)(rC)(rU)(rU)(rG)(rA)(rA)(rA)(rA)(rA)(rG)(rU)(rG)
(rG)(rC)(rA)(rC)(rC)(rG)(rA)(rG)(rU)(rC)(rG)(rG)(rU)(rG)(rC)(rU)(rU)(rU)(rU)(rU) - 3′

Synthesized by conjugation of:

5′ - (rG)(rG)(rG)(rC)(rG)(rA)(rG)(rG)(rA)(rG)(rC)(**rA**)(rG)(rU)(rU)(rC)(rA)(rC)(rC)
(rG)(rG)(rU)(rU)(rU)(rU)(rA)(rG)(rA)(rG)(rC)(rU)-**Tz** - 3′

and

5′ - **TCO**-(rA)(rG)(rA)(rA)(rA)(rU)(rA)(rG)(rC)(rA)(rA) (rG)(rU)(rU)(rA)(rA)(rA)(rA)(rU)(rA)
(rA)(rG)(rG)(rC)(rU)(rA)(rG)(rU)(rC)(rC)(rG)(rU)(rU)(rA)(rU)(rC)(rA)(rA)(rC)(rU)(rU)(rG)
(rA)(rA)(rA)(rA)(rA)(rG)(rU)(rG)(rG)(rC)(rA)(rC)(rC)(rG)(rA)(rG)(rU)(rC)(rG)(rG)(rU)(rG)
(rC)(rU)(rU)(rU)(rU)(rU) - 3′

eGFP-N1plasmid targeting **sgRNA 6** containing three U to A mismatches

5′ - (rG)(rG)(rG)(rC)(rG)(rA)(rG)(rG)(rA)(rG)(rC)(**rA**)(rG)(**rA**)(**rA**)(rC)(rA)(rC)(rC)
(rG)(rG)(rU)(rU)(rU)(rU)(rA)(rG)(rA)(rG)(rC)(rU)-**Tz-TCO**-(rA)(rG)(rA)(rA)(rA)(rU)(rA)
(rG)(rC)(rA)(rA) (rG)(rU)(rU)(rA)(rA)(rA)(rA)(rU)(rA)(rA)(rG)(rG)(rC)(rU)(rA)(rG)(rU)
(rC)(rC)(rG)(rU)(rU)(rA)(rU)(rC)(rA)(rA)(rC)(rU)(rU)(rG)(rA)(rA)(rA)(rA)(rA)(rG)(rU)(rG)
(rG)(rC)(rA)(rC)(rC)(rG)(rA)(rG)(rU)(rC)(rG)(rG)(rU)(rG)(rC)(rU)(rU)(rU)(rU)(rU) - 3′

Synthesized by conjugation of:

5′ - (rG)(rG)(rG)(rC)(rG)(rA)(rG)(rG)(rA)(rG)(rC)(**rA**)(rG)(**rA**)(**rA**)(rC)(rA)(rC)(rC)
(rG)(rG)(rU)(rU)(rU)(rU)(rA)(rG)(rA)(rG)(rC)(rU)-**Tz** - 3′

and

5′ - **TCO**-(rA)(rG)(rA)(rA)(rA)(rU)(rA)(rG)(rC)(rA)(rA) (rG)(rU)(rU)(rA)(rA)(rA)(rA)(rU)(rA)
(rA)(rG)(rG)(rC)(rU)(rA)(rG)(rU)(rC)(rC)(rG)(rU)(rU)(rA)(rU)(rC)(rA)(rA)(rC)(rU)(rU)(rG)
(rA)(rA)(rA)(rA)(rA)(rG)(rU)(rG)(rG)(rC)(rA)(rC)(rC)(rG)(rA)(rG)(rU)(rC)(rG)(rG)(rU)(rG)
(rC)(rU)(rU)(rU)(rU)(rU) - 3′

eGFP-N1plasmid targeting **sgRNA 7** containing three $m^1$A modifications

5′ - (rG)(rG)(rG)(rC)(rG)(**$m^1$A**)(rG)(rG)(**$m^1$A**)(rG)(rC)(rU)(rG)(rU)(rU)(rC)(**$m^1$A**)(rC)
(rC)(rG)(rG)(rU)(rU)(rU)(rU)(rA)(rG)(rA)(rG)(rC)(rU)-**Tz-TCO**-(rA)(rG)(rA)(rA)(rA)(rU)(rA)
(rG)(rC)(rA)(rA) (rG)(rU)(rU)(rA)(rA)(rA)(rA)(rU)(rA)(rA)(rG)(rG)(rC)(rU)(rA)(rG)(rU)
(rC)(rC)(rG)(rU)(rU)(rA)(rU)(rC)(rA)(rA)(rC)(rU)(rU)(rG)(rA)(rA)(rA)(rA)(rA)(rG)(rU)(rG)
(rG)(rC)(rA)(rC)(rC)(rG)(rA)(rG)(rU)(rC)(rG)(rG)(rU)(rG)(rC)(rU)(rU)(rU)(rU)(rU) - 3′

Synthesized by conjugation of:

5′ - (rG)(rG)(rG)(rC)(rG)(**$m^1$A**)(rG)(rG)(**$m^1$A**)(rG)(rC)(rU)(rG)(rU)(rU)(rC)(**$m^1$A**)(rC)
(rC)(rG)(rG)(rU)(rU)(rU)(rU)(rA)(rG)(rA)(rG)(rC)(rU)-**Tz** - 3′

and

5′ - **TCO**-(rA)(rG)(rA)(rA)(rA)(rU)(rA)(rG)(rC)(rA)(rA) (rG)(rU)(rU)(rA)(rA)(rA)(rA)(rU)(rA)
(rA)(rG)(rG)(rC)(rU)(rA)(rG)(rU)(rC)(rC)(rG)(rU)(rU)(rA)(rU)(rC)(rA)(rA)(rC)(rU)(rU)(rG)
(rA)(rA)(rA)(rA)(rA)(rG)(rU)(rG)(rG)(rC)(rA)(rC)(rC)(rG)(rA)(rG)(rU)(rC)(rG)(rG)(rU)(rG)
(rC)(rU)(rU)(rU)(rU)(rU) - 3′

eGFP-N1plasmid targeting **sgRNA 8** containing three $m^6$A modifications

5′ - (rG)(rG)(rG)(rC)(rG)(**$m^6$A**)(rG)(rG)(**$m^6$A**)(rG)(rC)(rU)(rG)(rU)(rU)(rC)(**$m^6$A**)(rC)
(rC)(rG)(rG)(rU)(rU)(rU)(rU)(rA)(rG)(rA)(rG)(rC)(rU)-**Tz-TCO**-(rA)(rG)(rA)(rA)(rA)(rU)(rA)
(rG)(rC)(rA)(rA) (rG)(rU)(rU)(rA)(rA)(rA)(rA)(rU)(rA)(rA)(rG)(rG)(rC)(rU)(rA)(rG)(rU)
(rC)(rC)(rG)(rU)(rU)(rA)(rU)(rC)(rA)(rA)(rC)(rU)(rU)(rG)(rA)(rA)(rA)(rA)(rA)(rG)(rU)(rG)
(rG)(rC)(rA)(rC)(rC)(rG)(rA)(rG)(rU)(rC)(rG)(rG)(rU)(rG)(rC)(rU)(rU)(rU)(rU)(rU) - 3′

Synthesized by conjugation of:

5′ - (rG)(rG)(rG)(rC)(rG)(**$m^6$A**)(rG)(rG)(**$m^6$A**)(rG)(rC)(rU)(rG)(rU)(rU)(rC)(**$m^6$A**)(rC)
(rC)(rG)(rG)(rU)(rU)(rU)(rU)(rA)(rG)(rA)(rG)(rC)(rU)-**Tz** - 3′

and

5′ - **TCO**-(rA)(rG)(rA)(rA)(rA)(rU)(rA)(rG)(rC)(rA)(rA) (rG)(rU)(rU)(rA)(rA)(rA)(rA)(rU)(rA)
(rA)(rG)(rG)(rC)(rU)(rA)(rG)(rU)(rC)(rC)(rG)(rU)(rU)(rA)(rU)(rC)(rA)(rA)(rC)(rU)(rU)(rG)
(rA)(rA)(rA)(rA)(rA)(rG)(rU)(rG)(rG)(rC)(rA)(rC)(rC)(rG)(rA)(rG)(rU)(rC)(rG)(rG)(rU)(rG)
(rC)(rU)(rU)(rU)(rU)(rU) - 3′

eGFP-N1plasmid targeting **sgRNA 9** containing three $s^2U$ modifications

5′ - (rG)(rG)(rG)(rC)(rG)(rA)(rG)(rG)(rA)(rG)(rC)(**$s^2$U**)(rG)(**$s^2$U**)(**$s^2$U**)(rC)(rA)(rC)(rC)(rG)(rG)(rU)(rU)(rU)(rU)(rA)(rG)(rA)(rG)(rC)(rU)-**Tz-TCO**-(rA)(rG)(rA)(rA)(rA)(rU)(rA)(rG)(rC)(rA)(rA) (rG)(rU)(rU)(rA)(rA)(rA)(rA)(rU)(rA)(rA)(rG)(rG)(rC)(rU)(rA)(rG)(rU)(rC)(rC)(rG)(rU)(rU)(rA)(rU)(rC)(rA)(rA)(rC)(rU)(rU)(rG)(rA)(rA)(rA)(rA)(rA)(rG)(rU)(rG)(rG)(rC)(rA)(rC)(rC)(rG)(rA)(rG)(rU)(rC)(rG)(rG)(rU)(rG)(rC)(rU)(rU)(rU)(rU)(rU) - 3′

Synthesized by conjugation of:

5′ - (rG)(rG)(rG)(rC)(rG)(rA)(rG)(rG)(rA)(rG)(rC)(**$s^2$U**)(rG)(**$s^2$U**)(**$s^2$U**)(rC)(rA)(rC)(rC)(rG)(rG)(rU)(rU)(rU)(rU)(rA)(rG)(rA)(rG)(rC)(rU)-**Tz** - 3′

and

5′ - **TCO**-(rA)(rG)(rA)(rA)(rA)(rU)(rA)(rG)(rC)(rA)(rA) (rG)(rU)(rU)(rA)(rA)(rA)(rA)(rU)(rA)(rA)(rG)(rG)(rC)(rU)(rA)(rG)(rU)(rC)(rC)(rG)(rU)(rU)(rA)(rU)(rC)(rA)(rA)(rC)(rU)(rU)(rG)(rA)(rA)(rA)(rA)(rA)(rG)(rU)(rG)(rG)(rC)(rA)(rC)(rC)(rG)(rA)(rG)(rU)(rC)(rG)(rG)(rU)(rG)(rC)(rU)(rU)(rU)(rU)(rU) - 3′

eGFP-N1plasmid targeting **sgRNA 10** containing three $s^4U$ modifications

5′ - (rG)(rG)(rG)(rC)(rG)(rA)(rG)(rG)(rA)(rG)(rC)(**$s^4$U**)(rG)(**$s^4$U**)(**$s^4$U**)(rC)(rA)(rC)(rC)(rG)(rG)(rU)(rU)(rU)(rU)(rA)(rG)(rA)(rG)(rC)(rU)-**Tz-TCO**-(rA)(rG)(rA)(rA)(rA)(rU)(rA)(rG)(rC)(rA)(rA) (rG)(rU)(rU)(rA)(rA)(rA)(rA)(rU)(rA)(rA)(rG)(rG)(rC)(rU)(rA)(rG)(rU)(rC)(rC)(rG)(rU)(rU)(rA)(rU)(rC)(rA)(rA)(rC)(rU)(rU)(rG)(rA)(rA)(rA)(rA)(rA)(rG)(rU)(rG)(rG)(rC)(rA)(rC)(rC)(rG)(rA)(rG)(rU)(rC)(rG)(rG)(rU)(rG)(rC)(rU)(rU)(rU)(rU)(rU) - 3′

Synthesized by conjugation of:

5′ - (rG)(rG)(rG)(rC)(rG)(rA)(rG)(rG)(rA)(rG)(rC)(**$s^4$U**)(rG)(**$s^4$U**)(**$s^4$U**)(rC)(rA)(rC)(rC)(rG)(rG)(rU)(rU)(rU)(rU)(rA)(rG)(rA)(rG)(rC)(rU)-**Tz** - 3′

and

5′ - **TCO**-(rA)(rG)(rA)(rA)(rA)(rU)(rA)(rG)(rC)(rA)(rA) (rG)(rU)(rU)(rA)(rA)(rA)(rA)(rU)(rA)(rA)(rG)(rG)(rC)(rU)(rA)(rG)(rU)(rC)(rC)(rG)(rU)(rU)(rA)(rU)(rC)(rA)(rA)(rC)(rU)(rU)(rG)(rA)(rA)(rA)(rA)(rA)(rG)(rU)(rG)(rG)(rC)(rA)(rC)(rC)(rG)(rA)(rG)(rU)(rC)(rG)(rG)(rU)(rG)(rC)(rU)(rU)(rU)(rU)(rU) - 3′

| sgRNA | Oligo Code | Sequence (5′ - 3′) | Calculated MW (Da) | Observed MW (Da) |
|---|---|---|---|---|
| **sgRNA 1** | 25mer-Tz | (rG)(rG)(rG)(rC)(rG)(rC)(rU)(rU)(rG)(rU)(rU)(rU)(rC)(rG)(rG)(rC)(rG)(rU)(rG)(rG)(rG) (rU)(rA)(rG)-**Tz** | 8335.1 | 8336.5 |
| **sgRNA 2** | 28mer-Tz | (rG)(rG)(rG)(rC)(rG)(rC)(rU)(rU)(rG)(rU)(rU)(rU)(rC)(rG)(rG)(rC)(rG)(rU)(rG)(rG)(rG) (rU)(rA)(rG)(rU)(rU)(rU)(rU)-**Tz** | 9254.9 | 9254.8 |
| **sgRNA 3** | 34mer-Tz | (rG)(rG)(rG)(rC)(rG)(rC)(rU)(rU)(rG)(rU)(rU)(rU)(rC)(rG)(rG)(rC)(rG)(rU)(rG)(rG)(rG)(rU)(rA)(rG)(rU)(rU)(rU)(rU)(rA)(rG)(rA)(rG)(rC)(rU)-**Tz** | 11215.7 | 11215.8 |
| **sgRNA 4** | 31mer-Tz | (rG)(rG)(rG)(rC)(rG)(rA)(rG)(rG)(rA)(rG)(rC)(rU)(rG)(rU)(rU)(rC)(rA)(rC)(rC)(rG)(rG)(rU)(rU)(rU)(rU)(rA)(rG)(rA)(rG)(rC)(rU)-**Tz** | 10263.4 | 10263.6 |
| **sgRNA 5** | 31mer-Tz one U to A mismatch | (rG)(rG)(rG)(rC)(rG)(rA)(rG)(rG)(rA)(rG)(rC)(**rA**)(rG)(rU)(rU)(rC)(rA)(rC)(rC)(rG)(rG)(rU)(rU)(rU)(rU)(rA)(rG)(rA)(rG)(rC)(rU)-**Tz** | 10288.1 | 10288.2 |
| **sgRNA 6** | 31mer-Tz three U to A mismatch | (rG)(rG)(rG)(rC)(rG)(rA)(rG)(rG)(rA)(rG)(rC)(**rA**)(rG)(**rA**)(**rA**)(rC)(rA)(rC)(rC)(rG)(rG)(rU)(rU)(rU)(rU)(rA)(rG)(rA)(rG)(rC)(rU)-**Tz** | 10333.3 | 10333.5 |
| **sgRNA 7** | 31mer-Tz three $m^1A$ | (rG)(rG)(rG)(rC)(rG)(**$m^1A$**)(rG)(rG)(**$m^1A$**)(rG)(rC)(rU)(rG)(rU)(rU)(rC)(**$m^1A$**)(rC)(rC)(rG)(rG)(rU)(rU)(rU)(rU)(rA)(rG)(rA)(rG)(rC)(rU)-**Tz** | 10305.1 | 10305.3 |
| **sgRNA 8** | 31mer-Tz three $m^6A$ | (rG)(rG)(rG)(rC)(rG)(**$m^6A$**)(rG)(rG)(**$m^6A$**)(rG)(rC)(rU)(rG)(rU)(rU)(rC)(**$m^6A$**)(rC)(rC)(rG)(rG)(rU)(rU)(rU)(rU)(rA)(rG)(rA)(rG)(rC)(rU)-**Tz** | 10305.1 | 10304.5 |
| **sgRNA 9** | 31mer-Tz three $s^2U$ | (rG)(rG)(rG)(rC)(rG)(rA)(rG)(rG)(rA)(rG)(rC)(**$s^2U$**)(rG)(**$s^2U$**)(**$s^2U$**)(rC)(rA)(rC)(rC)(rG)(rG)(rU)(rU)(rU)(rU)(rA)(rG)(rA)(rG)(rC)(rU)-**Tz** | 10311.4 | 10311.0 |
| **sgRNA 10** | 31mer-Tz three $s^4U$ | (rG)(rG)(rG)(rC)(rG)(rA)(rG)(rG)(rA)(rG)(rC)(**$s^4U$**)(rG)(**$s^4U$**)(**$s^4U$**)(rC)(rA)(rC)(rC)(rG)(rG)(rU)(rU)(rU)(rU)(rA)(rG)(rA)(rG)(rC)(rU)-**Tz** | 10309.5 | 10301.4 |

**Table S1.** ESI-MS analysis of the reported RNA oligonucleotides.

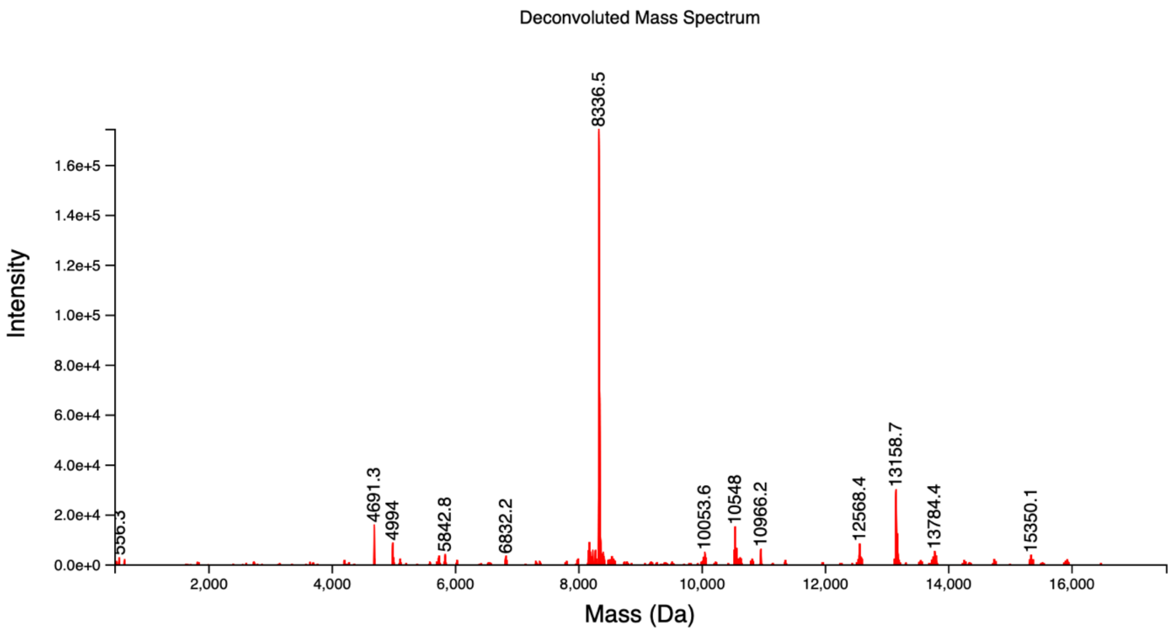


**Figure S1.** Deconvoluted ESI-MS spectrum of the 25mer-Tz that was used to form **sgRNA1**.

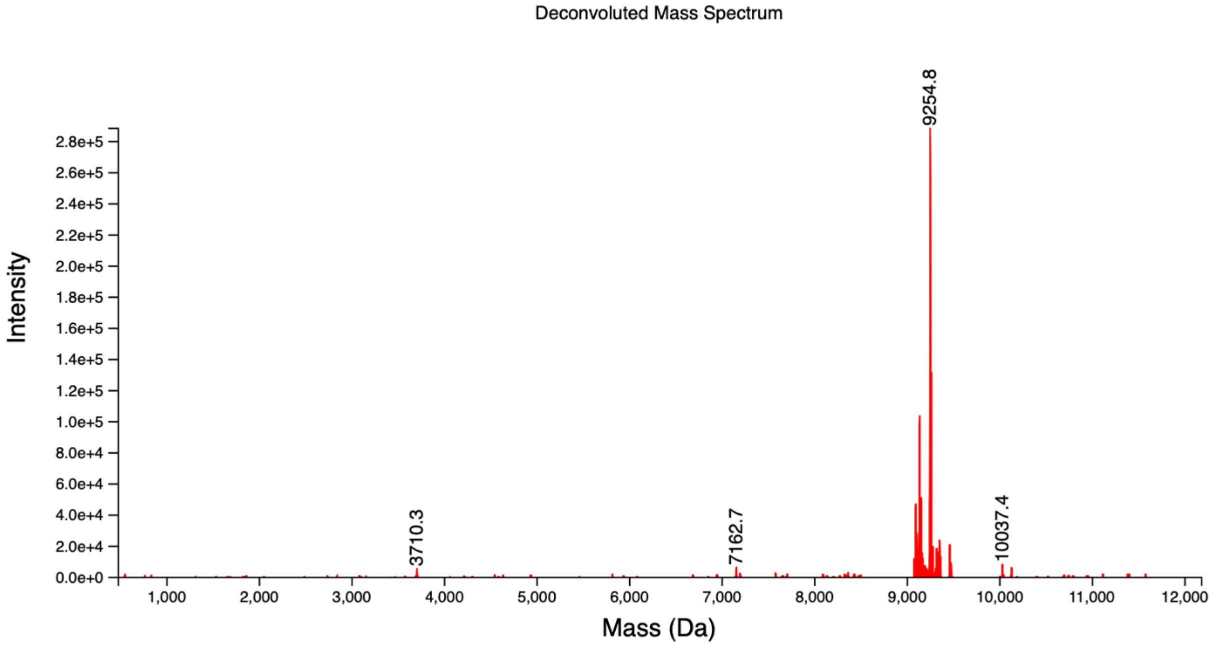


**Figure S2.** Deconvoluted ESI-MS spectrum of the 28mer-Tz that was used to form **sgRNA2**.

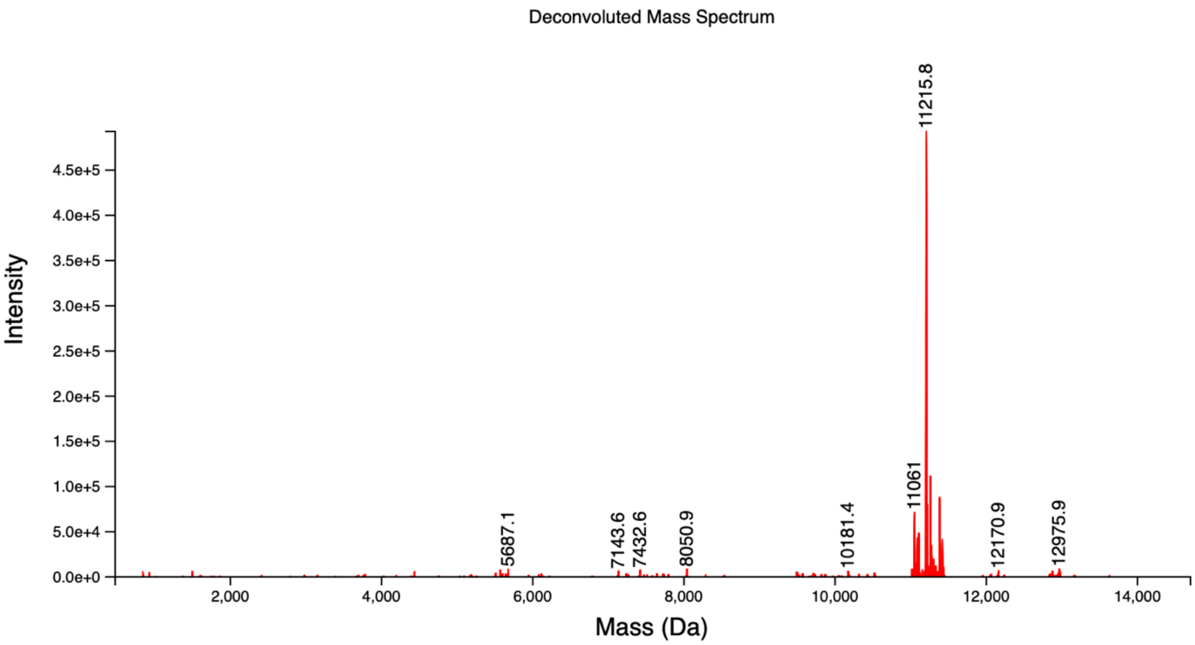


**Figure S3.** Deconvoluted ESI-MS spectrum of the 34mer-Tz that was used to form **sgRNA3**.

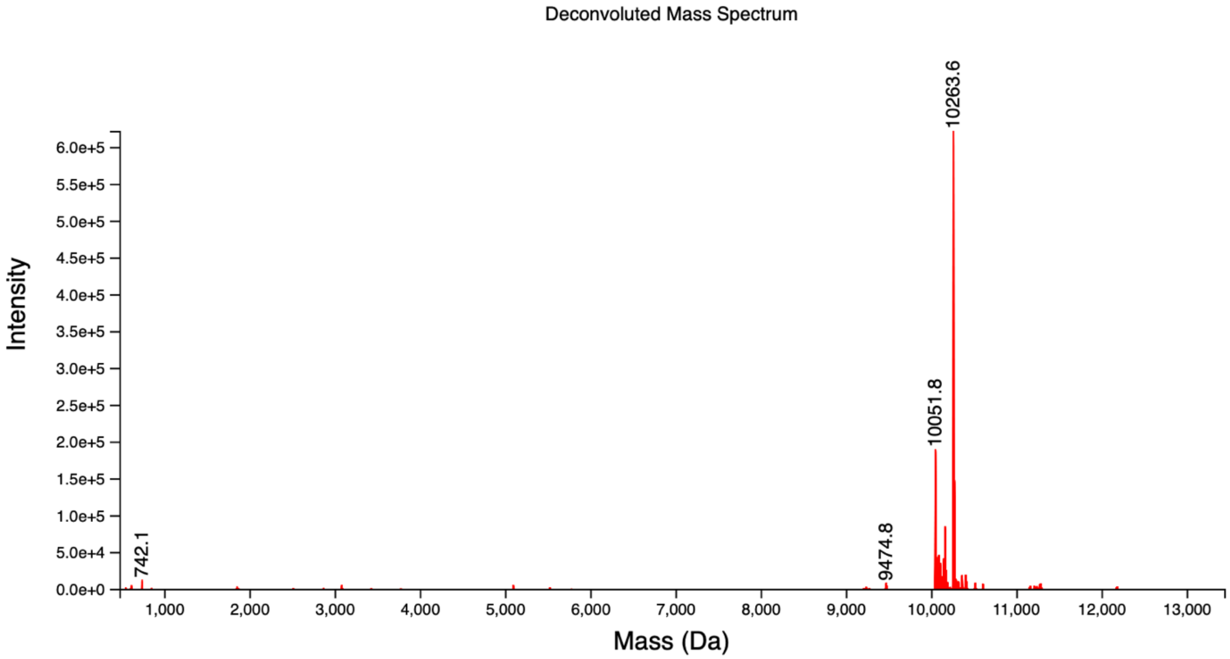


**Figure S4.** Deconvoluted ESI-MS spectrum of the 31mer-Tz that was used to form **sgRNA4**.

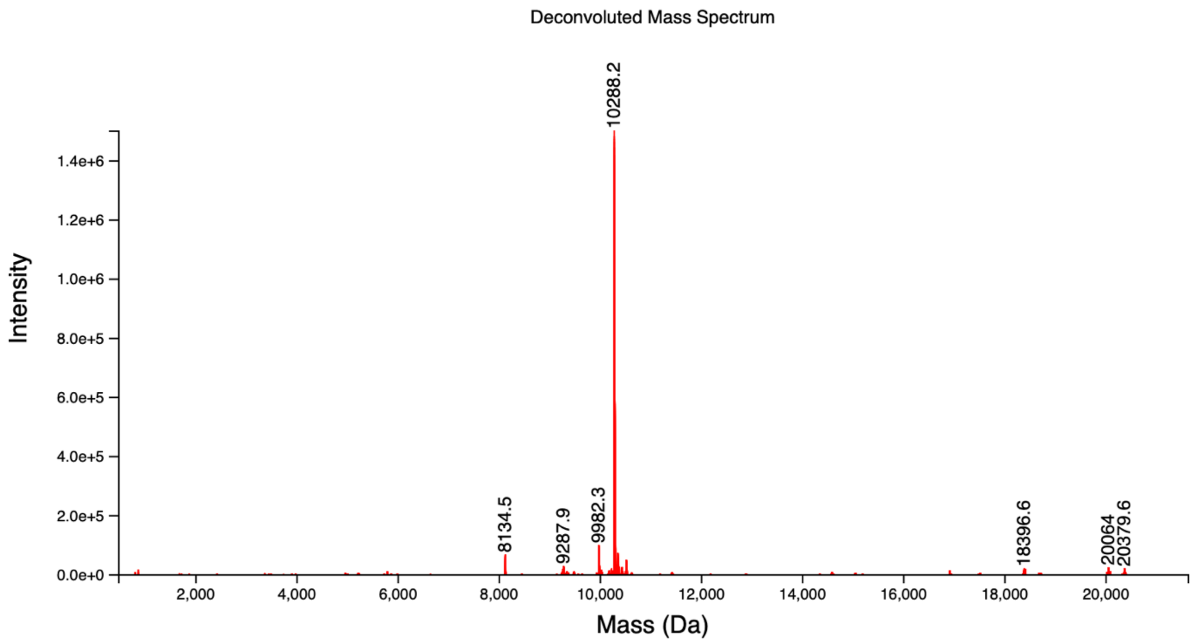


**Figure S5.** Deconvoluted ESI-MS spectrum of the 31mer-Tz that was used to form **sgRNA5**.

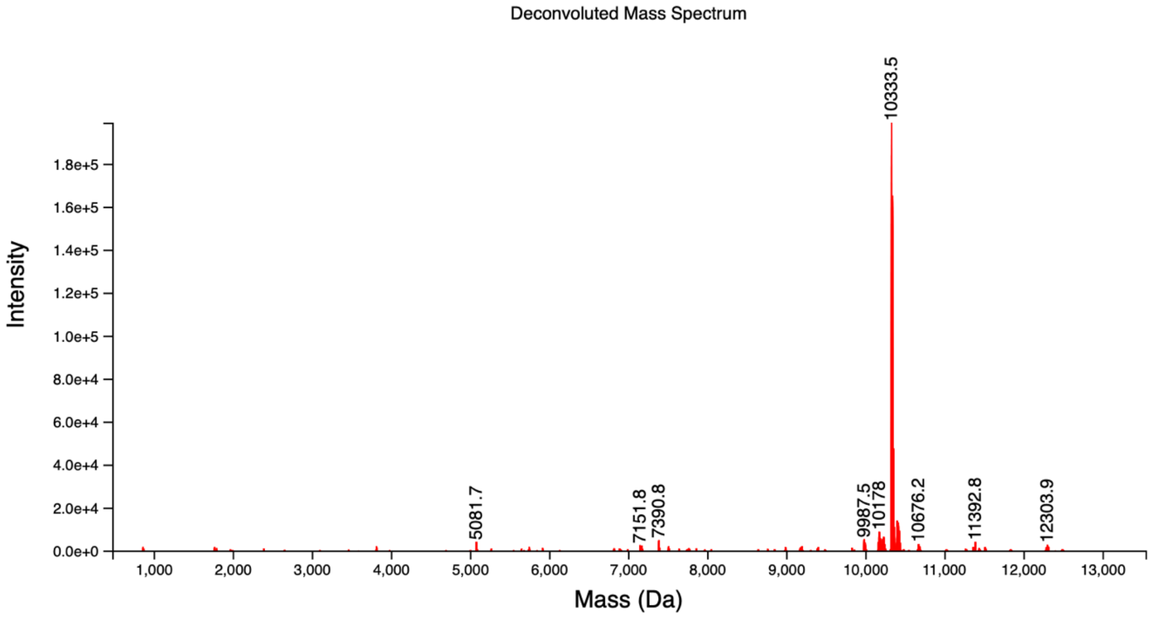


**Figure S6.** Deconvoluted ESI-MS spectrum of the 31mer-Tz that was used to form **sgRNA6**.

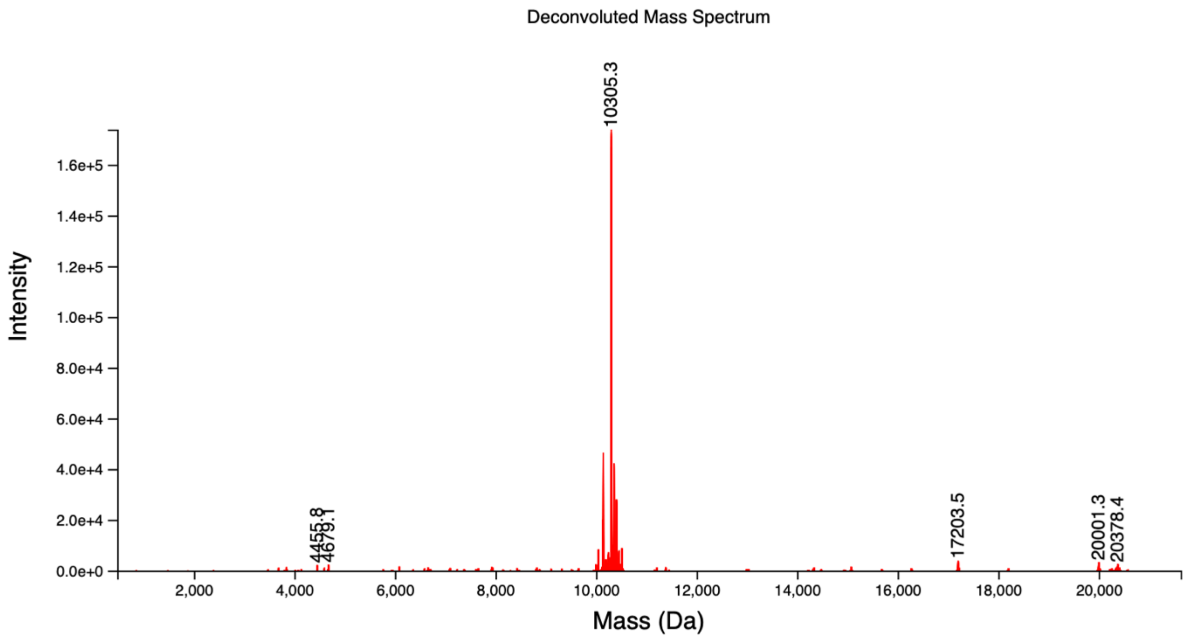


**Figure S7.** Deconvoluted ESI-MS spectrum of the 31mer-Tz that was used to form **sgRNA7**.

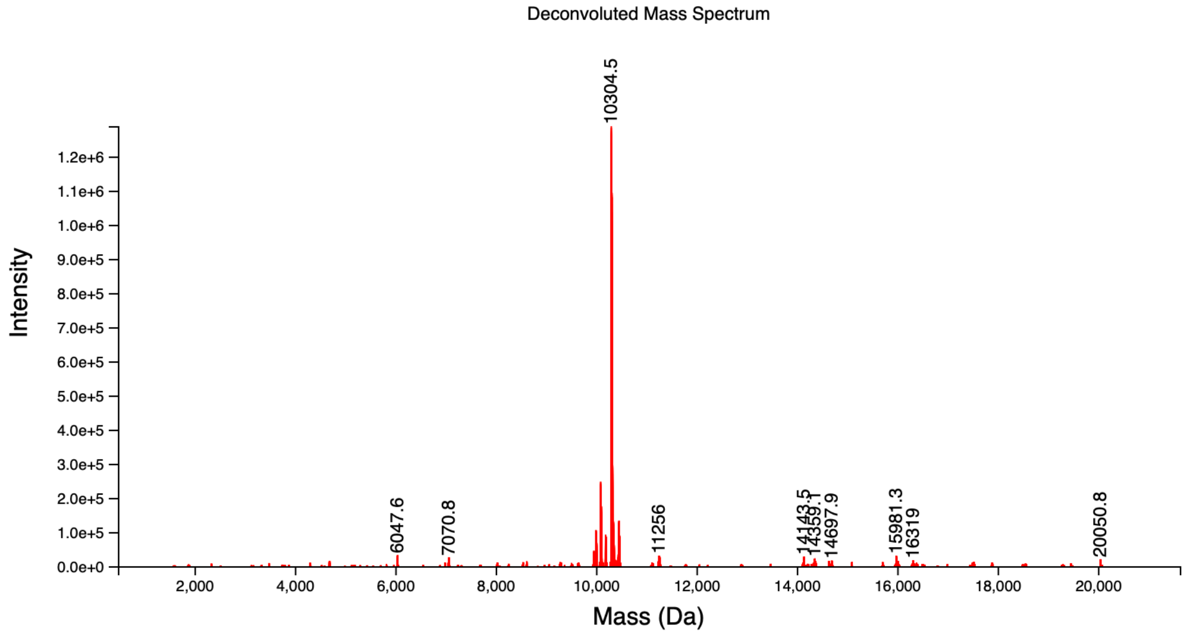


**Figure S8.** Deconvoluted ESI-MS spectrum of the 31mer-Tz that was used to form **sgRNA8**.

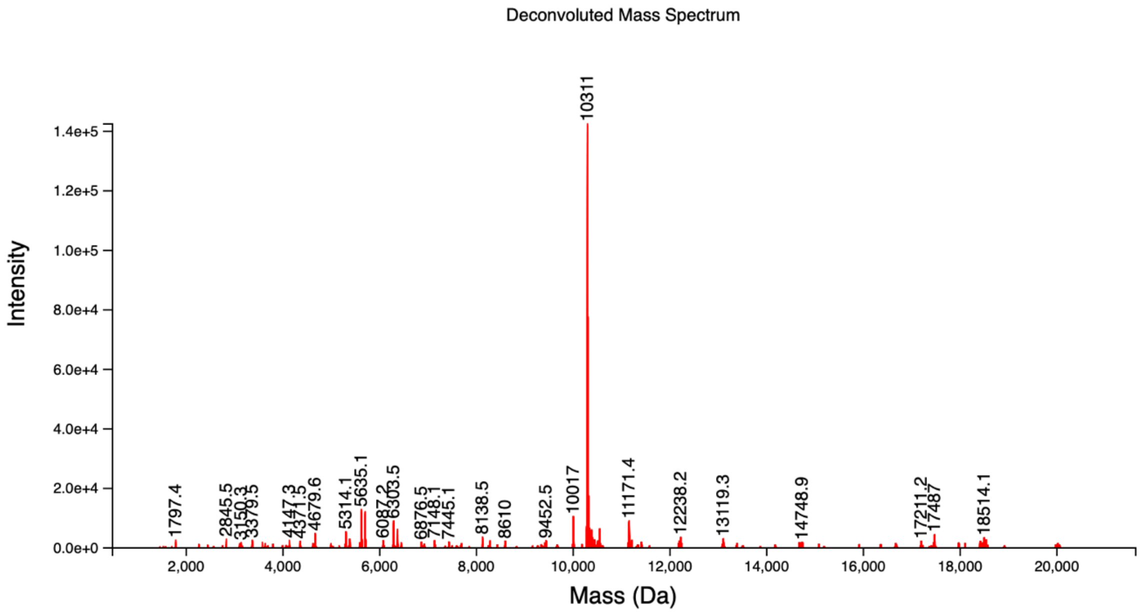


**Figure S9.** Deconvoluted ESI-MS spectrum of the 31mer-Tz that was used to form **sgRNA9**.

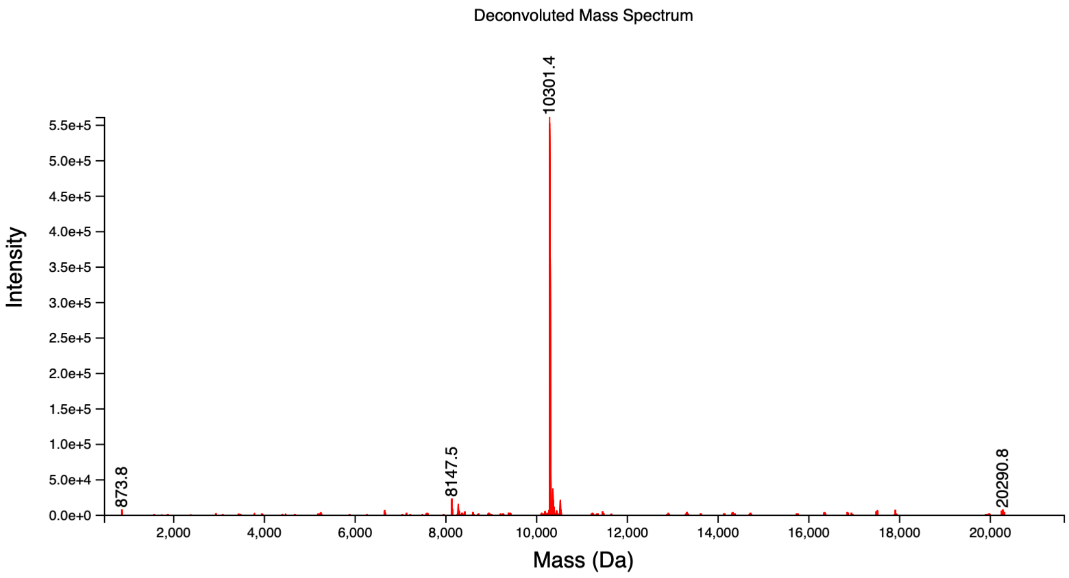


**Figure S10.** Deconvoluted ESI-MS spectrum of the 31mer-Tz that was used to form **sgRNA10**.

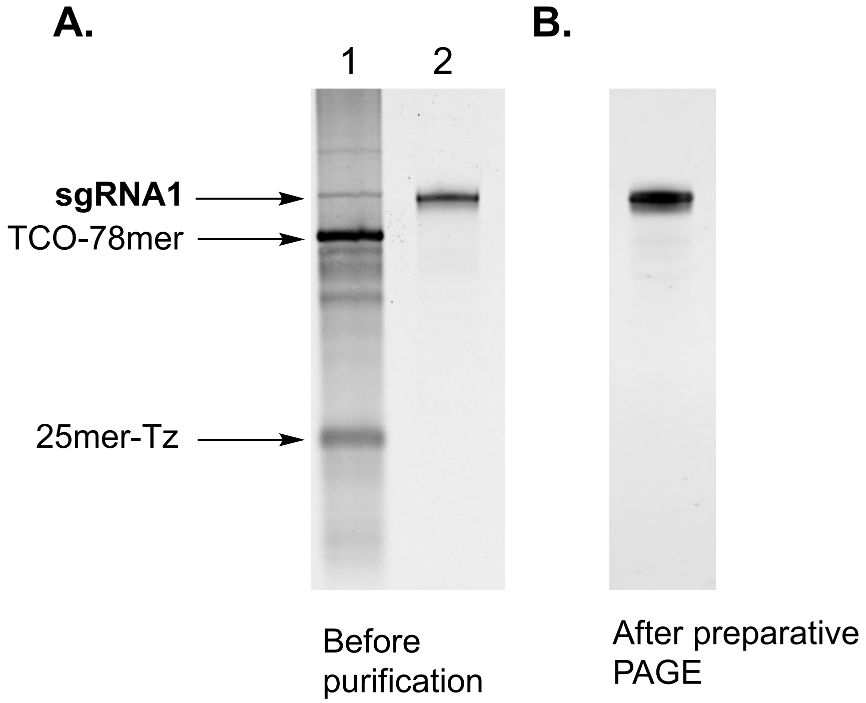


**Figure S11.** PAGE analysis of conjugation of 25mer-Tz and TCO-78mer. (**A.**) *Lane 1:* Crude reaction mixture, showing formation of **sgRNA1**; *Lane 2:* reference 103-nt long RNA. (**B.**) **sgRNA1** after purification by preparative PAGE.

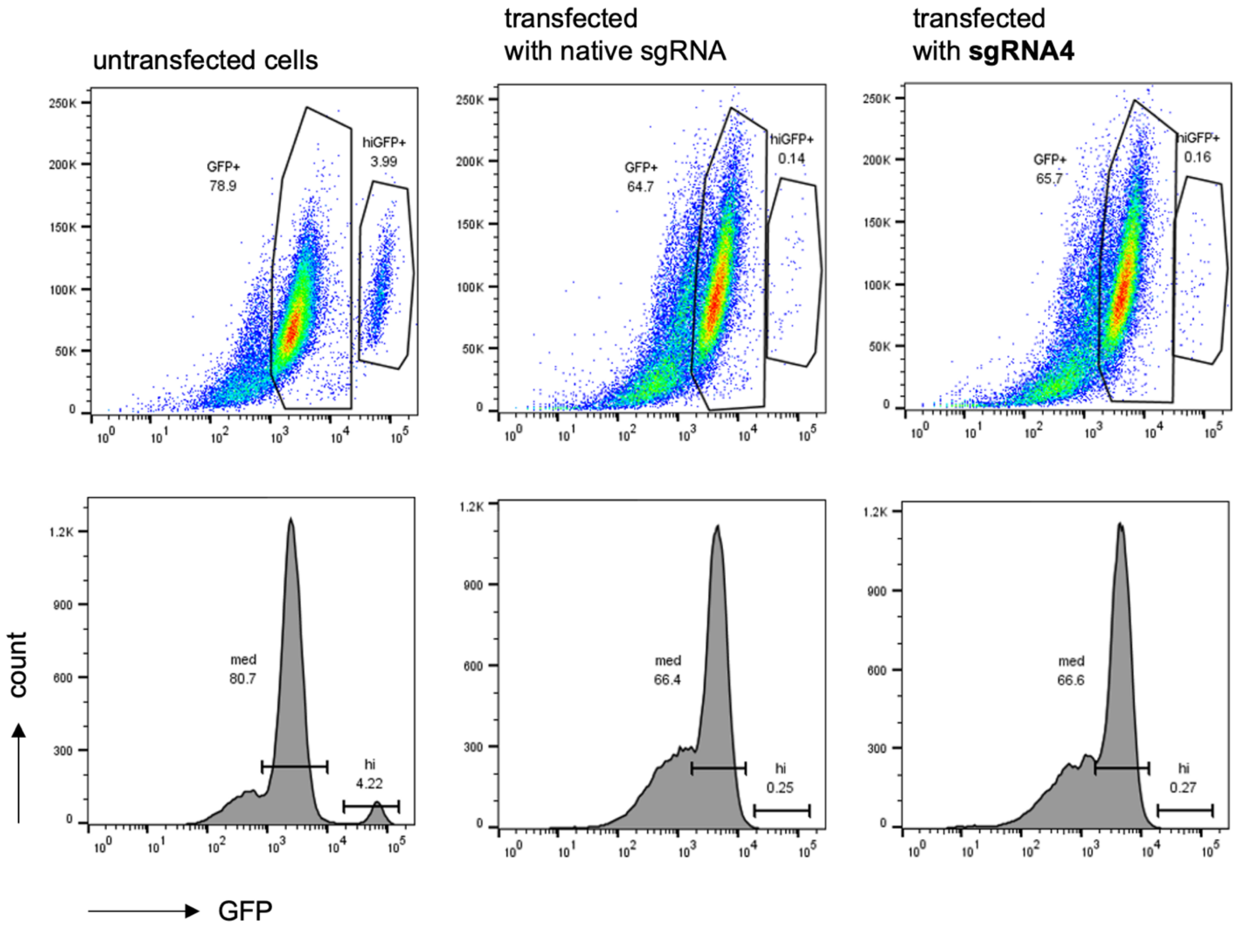


**Figure S12.** Flow cytometry analysis of GFP expression of GFP and Cas9-expressing HEK293T transfected with native sgRNA and **sgRNA4**.

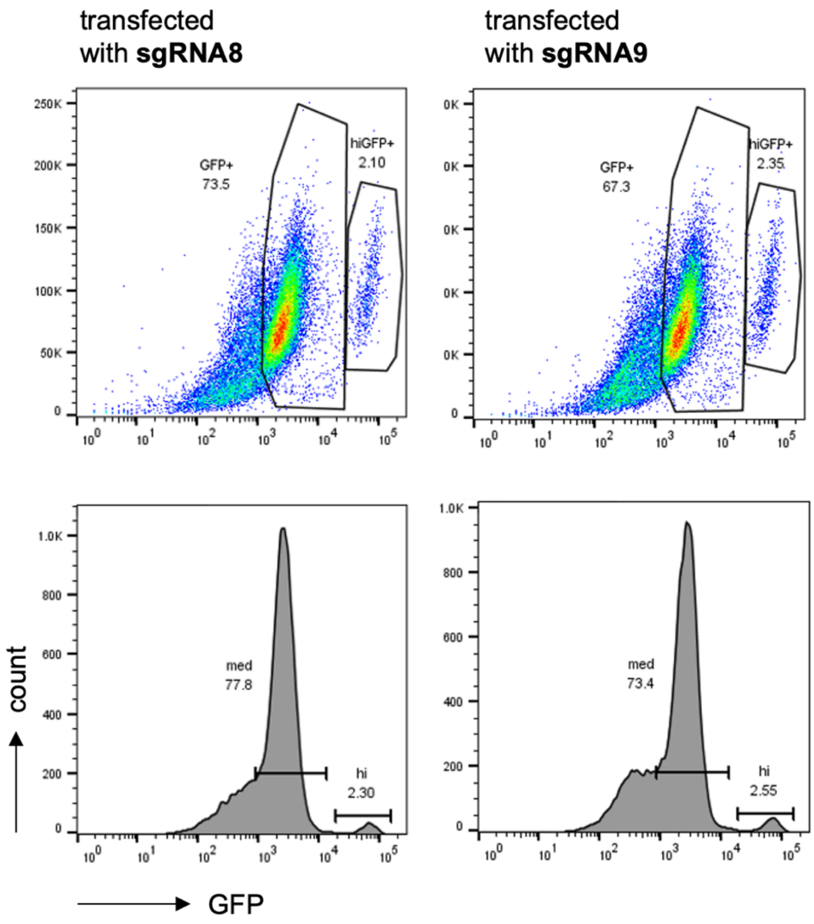


**Figure S13.** Flow cytometry analysis of GFP expression of GFP and Cas9-expressing HEK293T transfected with **sgRNA8** and **sgRNA9**.

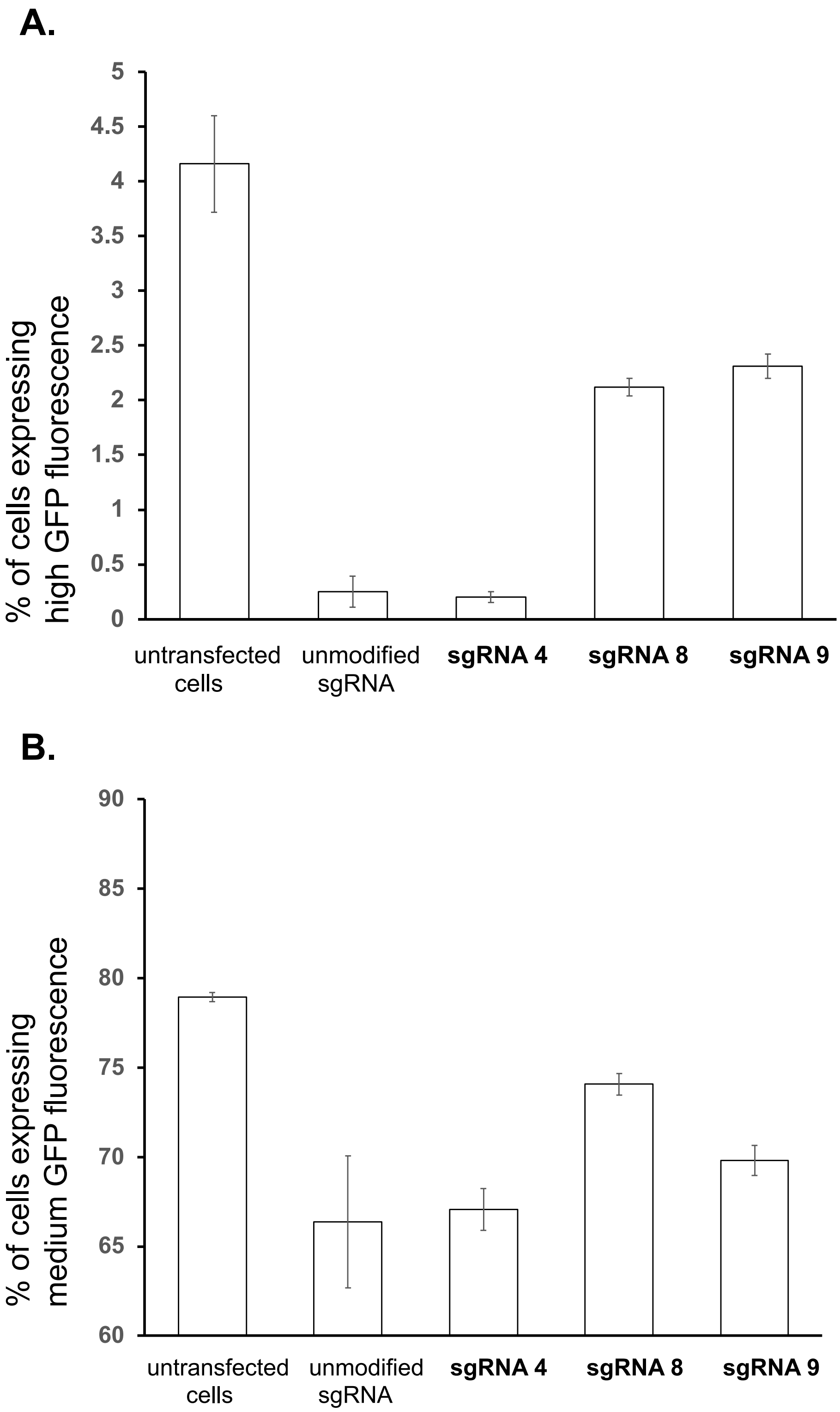


**Figure S14.** Flow cytometry analysis of GFP expression of GFP and Cas9-expressing HEK293T transfected with unmodified sgRNA, **sgRNA 4**, **sgRNA8** and **sgRNA9**. (**A.**) subpopulation of cells with high GFP fluorescence, (**B.**) subpopulation of cells with medium GFP fluorescence.

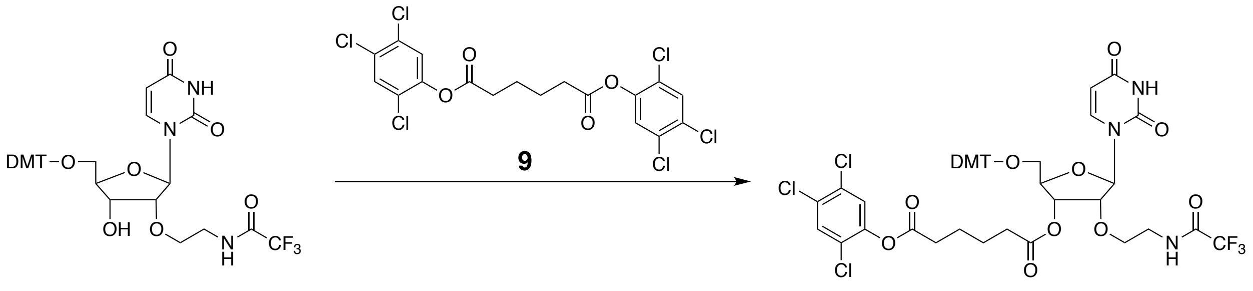


Compound **8** was synthesized following the previously reported procedure [Santner, T.; Hartl, M.; Bister, K.; Micura, R. Bioconjug. Chem. **2014**, *25*, 188-195]. Compound **9** (0.439 g, 0.87 mmol) was dissolved in a 1:1 solution of $CH_2Cl_2$/pyridine (8 mL) and a catalytic amount of 4-dimethylaminopyridine (~30 mg) was added. Compound **8** (0.200 g, 0.29 mmol) was co-evaporated with pyridine twice and dissolved in $CH_2Cl_2$ (2 mL). The solution was added dropwise over 1 h and stirred for 16 h at rt. The solution was directly purified using preparatory silica gel plate 2-5% MeOH/DCM to afford the title compound **10** as an off-white solid (0.116 g, 40%).

$^1$H NMR (500 MHz, $CDCl_3$) δ 7.95 (d, *J* = 8.1 Hz, 1H), 7.55 (s, 1H), 7.51 (s, 1H), 7.38 (d, *J* = 6.9 Hz, 2H), 7.35 – 7.23 (m, 8H), 6.87 (d, *J* = 8.5 Hz, 4H), 5.98 (s, 1H), 5.41 (d, *J* = 8.1 Hz, 1H), 5.20 (s, 1H), 4.33 – 4.24 (m, 2H), 3.81 (s, 6H), 3.67 (d, *J* = 9.9 Hz, 2H), 3.64 – 3.40 (m, 4H), 2.91 (q, *J* = 7.1 Hz, 1H), 2.67 (t, *J* = 6.1 Hz, 2H), 2.45 (d, *J* = 6.0 Hz, 2H), 1.80 (dd, *J* = 15.4, 6.5 Hz, 3H), 1.68 (s, 1H), 1.28 (s, 1H), 1.23 (t, *J* = 7.2 Hz, 2H).
$^{13}$C NMR (126 MHz, $CDCl_3$) δ 172.43 (s), 170.17 (s), 163.14 (s), 158.77 (d, *J* = 2.0 Hz), 151.05 (s), 145.74 (s), 143.96 (s), 139.35 (s), 134.83 (d, *J* = 11.0 Hz), 131.40 (s), 130.93 (s), 130.47 (s), 130.01 (t, *J* = 7.7 Hz), 128.00 (d, *J* = 1.7 Hz), 127.22 (s), 126.04 (s), 125.27 (s), 118.75 (s), 113.31 (s), 102.87 (s), 88.14 (s), 87.38 (s), 81.53 (s), 80.99 (s), 70.09 (s), 69.08 (s), 55.17 (s), 45.58 (s), 39.40 (s), 33.29 (d, *J* = 12.7 Hz), 24.21 – 23.85 (m), 9.29 (s).
HRMS (ESI) Calc'd for $C_{46}H_{43}Cl_3F_3N_3NaO_{12}$ $[M+Na]^+$ 1014.1762; found 1014.1760

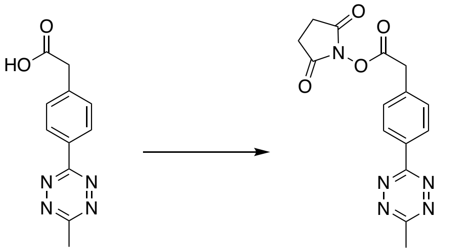


Compound **11** was synthesized following the previously described procedure [Yang, J.; Karver, M. R.; Li, W.; S., Swagat; Devaraj, N. K. Angew Chem. Int. Ed. **2012**, 51, 5222-5225]. Compound **11** (0.100 g, 0.434 mmol) and *N,N*-disuccinimidyl cabonate (0.222 g, 0.868 mmol) were dissolved in $CH_3CN$ (5 mL). DIPEA (0.150 mL, 0.869 mmol) was added dropwise and the reaction mixture was stirred for 4 h at rt. Quenched the reaction with water (1 mL) and extracted the product with EtOAc (2x50 mL). Dried the organic layers with $Na_2SO_4$ and removed solvents under reduced pressure. The crude product was coupled to RNA without further purification. Yield: 0.142 g (80%)

$^1$H NMR (500 MHz, $CDCl_3$) δ 8.63 (d, *J* = 8.2 Hz, 2H), 7.61 (d, *J* = 8.2 Hz, 2H), 4.08 (s, 2H), 3.13 (s, 5H), 2.88 (s, 7H).
HRMS (ESI) Calc'd for $C_{15}H_{14}N_5O_4$ $[M+1]^+$ 328.1040; found 328.1041

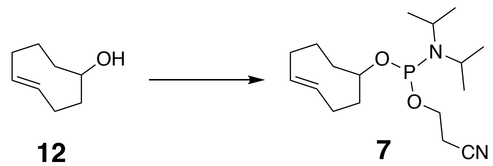

12 7

The compound **7** was synthesized using previously described procedure [Schoch, J.; Staudt, M.; Samanta, A.; Wiessler, M.; Jaeschke, A. Bioconjug. Chem. 2012, 23, 1382-1386.] The major diastereomer of *trans*-cyclooctenol (200 mg, 1.58 mmol) was dissolved in anhydrous $CH_2Cl_2$ (2 mL). DIPEA (0.82 ml, 4.75 mmol) was added dropwise. The mixture was cooled to 0°C and 2-cyanoethyl-N,N-diisopropylchlorophosphoramidite (0.38 mL, 1.74 mmol) was added dropwise. The reaction mixture was stirred at rt for 1 h. The reaction mixture was loaded directly onto a silica column. The title product was purified by flash chromatography using a gradient of EtOAc in Hexanes (0-20 %). Yield = 287 mg (56%)

$^1$H NMR (500 MHz, $CDCl_3$) δ 5.67 – 5.37 (m, 2H), 3.89 – 3.72 (m, 2H), 3.67 – 3.52 (m, 3H), 2.72 – 2.59 (m, 2H), 2.47 – 1.38 (m, 12H), 1.28 – 1.15 (m, 13H).
$^{31}$P NMR (121 MHz, $CDCl_3$): δ=146.1 145.5 (mixture of 2 diastereomers).
HRMS (ESI) Calc'd for $C_{17}H_{32}N_2O_2P$ $[M+1]^+$ 327.2196; found 327.2193

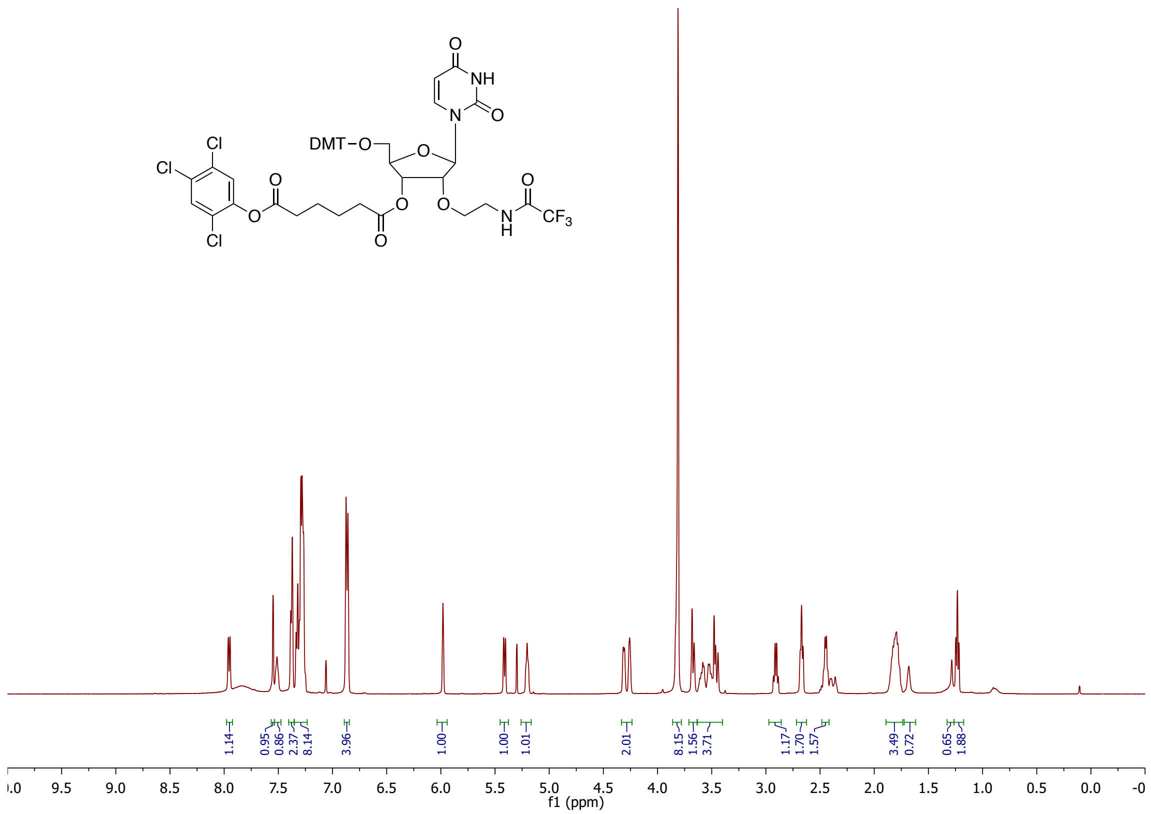
DMT–O
NH
N
O
Cl
CF3
H
1.14
0.95
0.86
2.37
8.14
3.96
1.00
1.00
1.01
2.01
8.15
1.56
3.71
1.17
1.70
1.57
3.49
0.72
0.65
1.88
f1 (ppm)

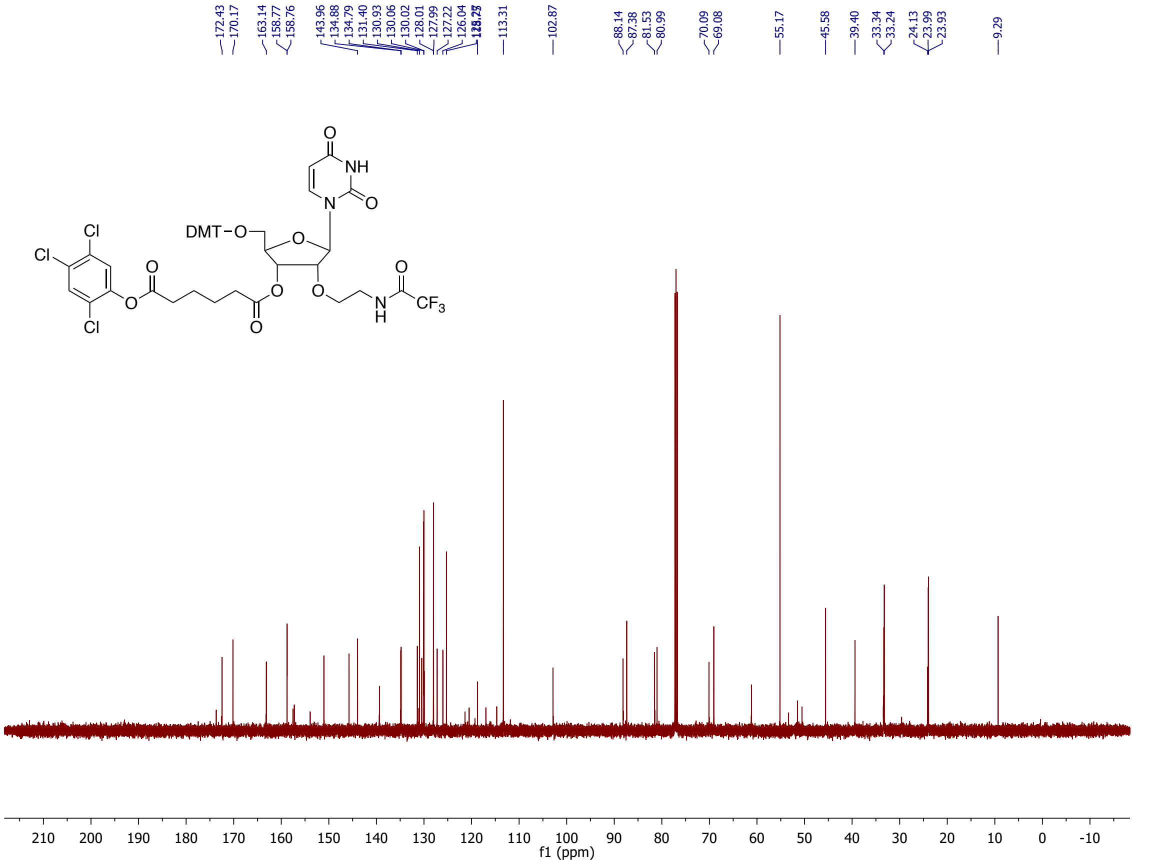
172.43
170.17
163.14
158.77
158.76
143.96
134.88
134.79
131.40
130.93
130.06
130.02
128.01
127.99
127.22
126.04
113.31
102.87
88.14
87.38
81.53
80.99
70.09
69.08
55.17
45.58
39.40
33.34
33.24
24.13
23.99
23.93
9.29
DMT–O
NH
N
O
Cl
CF3
H
f1 (ppm)

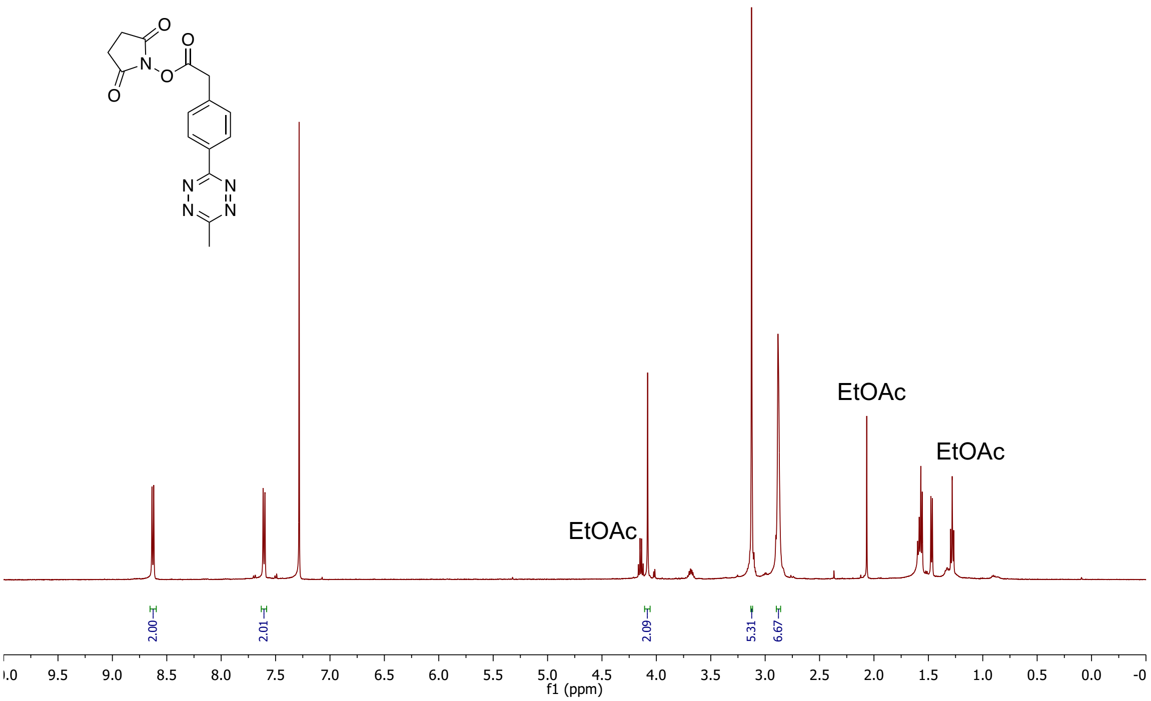
EtOAc
EtOAc
EtOAc
2.00
2.01
2.09
5.31
6.67
9.5
9.0
8.5
8.0
7.5
7.0
6.5
6.0
5.5
5.0
4.5
4.0
3.5
3.0
2.5
2.0
1.5
1.0
0.5
0.0
f1 (ppm)

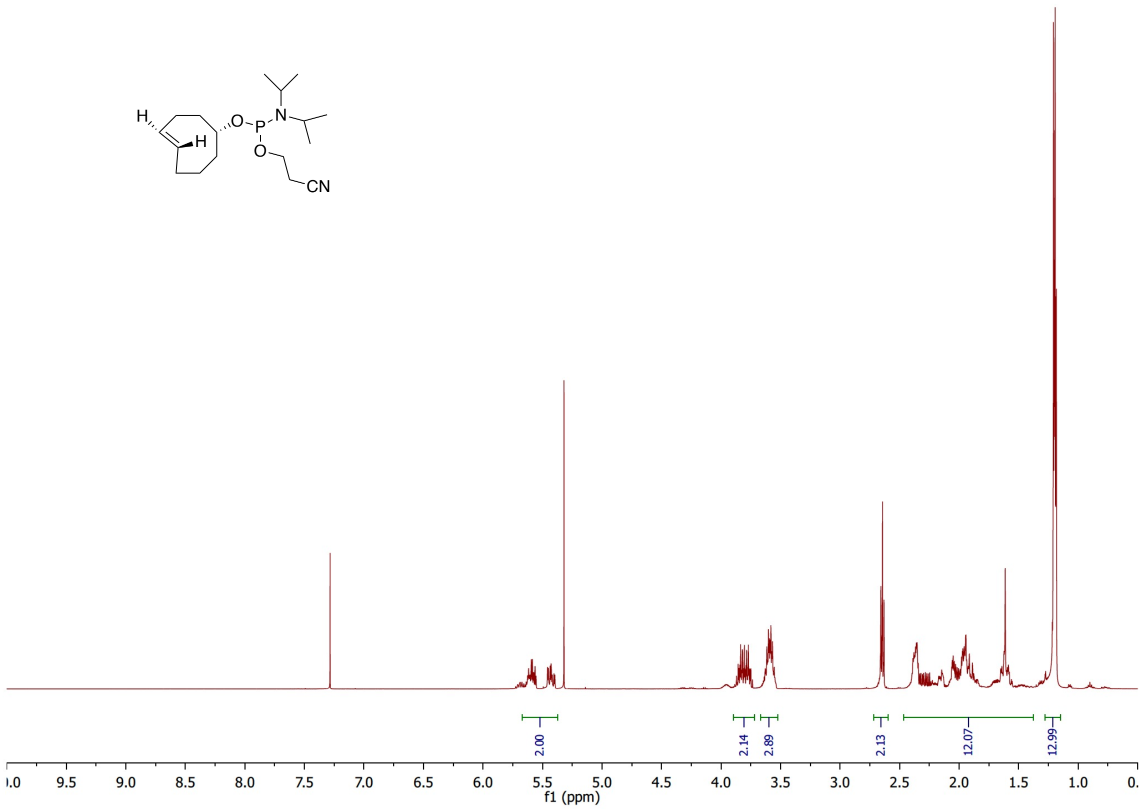
H
H
O
P
N
O
CN
2.00
2.14
2.89
2.13
12.07
12.99
f1 (ppm)

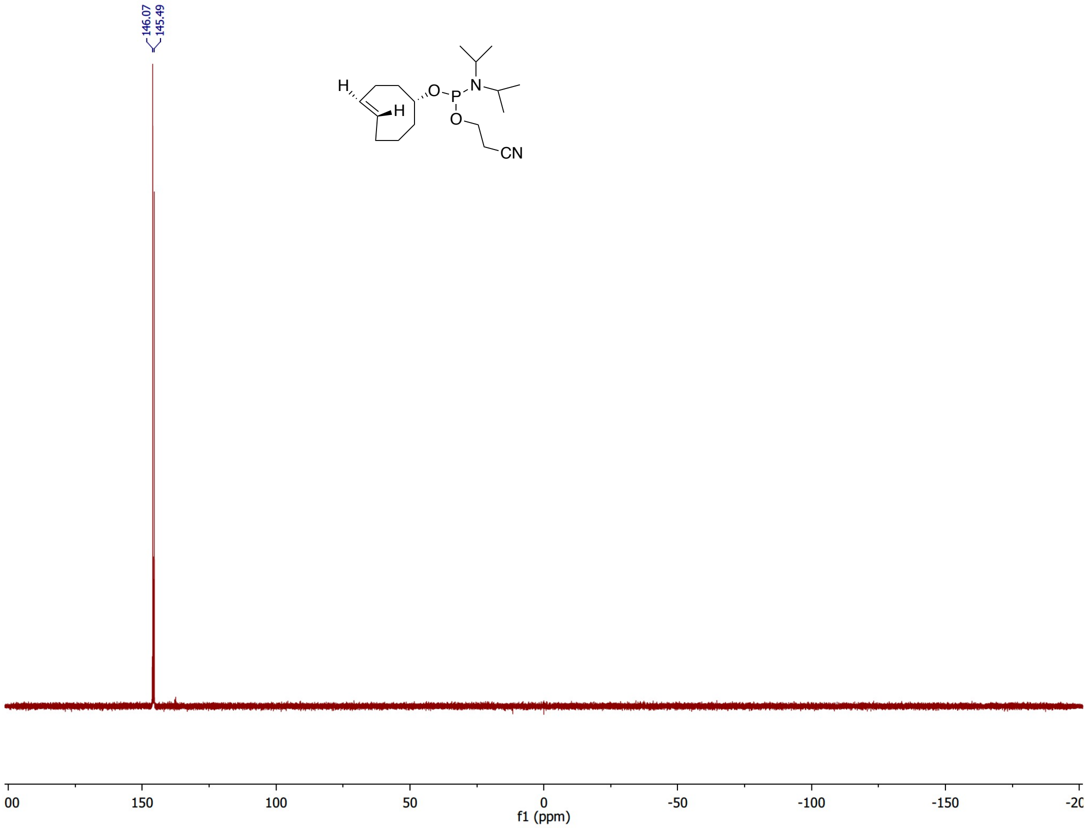
146.07
145.49
H
H
O
P
N
O
CN
f1 (ppm)
